# *home*RNA: A self-sampling kit for the collection of peripheral blood and stabilization of RNA

**DOI:** 10.1101/2021.02.08.430337

**Authors:** Amanda J. Haack, Fang Yun Lim, Dakota S. Kennedy, John H. Day, Karen N. Adams, Jing J. Lee, Erwin Berthier, Ashleigh B. Theberge

**Affiliations:** Department of Chemistry, University of Washington, Seattle, WA, USA, 98195; School of Medicine, University of Washington, Seattle, WA, USA, 98195; Institute of Translational Health Sciences, School of Medicine, University of Washington, Seattle, WA, USA, 98195; Department of Urology, School of Medicine, University of Washington, Seattle, WA, USA, 98195

## Abstract

Gene expression analysis (e.g., targeted gene panels, transcriptomics) from whole blood can elucidate mechanisms of immune function and aid in the discovery of biomarkers. Conventional venipuncture offers only a small snapshot of our broad immune landscape as immune responses may occur outside of the time and location parameters available for conventional venipuncture. A self-operated method that enables flexible sampling of liquid whole blood coupled with immediate stabilization of cellular RNA is instrumental in facilitating capture and preservation of acute or transient immune fluxes. To this end, we developed *home*RNA: a kit for self-collection of peripheral blood (∼0.5 mL) and immediate stabilization of cellular RNA, using the Tasso- SST™ blood collection device with a specially designed stabilizer tube containing RNA*later*™. To assess the feasibility of *home*RNA for self-collection and stabilization of whole blood RNA, we conducted a pilot study (n = 47 participants) where we sent *home*RNA to participants aged 21-69, located across 10 US states (94% successful blood collections, n = 61/65). Among participants who successfully collected blood, 93% reported no or minimal pain/discomfort using the kit (n = 39/42), and 79% reported very easy/somewhat easy stabilization protocol (n = 33/42). Total RNA yield from the stabilized samples ranged between 0.20 µg and 5.99 µg (mean = 1.51 µg), and all but one RNA Integrity Number (RIN) values were above 7.0 (mean = 8.1), indicating limited RNA degradation. Results from this study demonstrate the self-collection and RNA stabilization of whole blood with *home*RNA by participants themselves, in their own home.

## INTRODUCTION

Remote and contact-free laboratory testing is rapidly emerging as the new standard in patient care and clinical research, especially in light of the COVID-19 pandemic. However, blood sample collection remains a challenging procedure to perform remotely as venipuncture is resource-intensive, physically uncomfortable, and inflexible in regard to collection time and location.^1, 2^ Remote self-administered blood collection, on the other hand, offers many practical advantages, including 1) expanded lab testing for rural and remote medicine applications (i.e., telemedicine), 2) convenience for clinical research studies as well as the ability to recruit participants that are not able to come to the clinic (due to work schedules, caregiver responsibilities, mobility challenges, etc.), 3) the ability to capture acute and transient biomarker fluxes (e.g., immediately following an acute exposure, an asthma attack, or a flare in an autoimmune disease), and 4) opportunities to conduct longitudinal research studies that require frequent sample collections from the same individual over a short time course (e.g., daily blood collections); to-date these applications have been limited due to the logistical challenges associated with in-person venipuncture. Here, we will describe *home*RNA: a new technology to enable at-home collection and stabilization of whole blood cellular RNA for gene expression analysis.

An important example of an existing technology aimed at remote blood sampling is the use of dried blood spot (DBS) sampling. In DBS sampling, a lancet-based finger prick is used to draw blood, which is applied to a sampling paper and left to dry. The sampling paper containing the DBS is then mailed back to the lab for analysis.^3^ This technology has been applied to a variety of applications, including diagnostics and screening,^4–9^ therapeutic drug monitoring,^1, 10, 11^and other mechanistic biomolecule analysis.^12, 13^ Due to the increased use of DBS and convenience for remote sampling, tremendous research and development have been undertaken to improve the consistency and analysis of DBS samples.^5, 14–16^ In comparison to a dried blood spot, a liquid blood sample can provide a larger volume of blood (>100 µL). An increased sample volume may be desirable for applications such as genomics, transcriptomics, or the detection of rare analytes. Further, if adequately stabilized, liquid samples may provide a greater quantity and better quality of the desired analyte than DBS samples, such as a higher yield of minimally degraded total whole blood RNA.^17^ Another burgeoning class of blood sample collection devices is lancet-based devices that collect from capillary beds in the arm. The user activates the lancet by pushing a button, which then causes blood to flow into a collection receptacle.^18–21^ The Tasso-SST™ blood collection device used in this study falls under this category of sampling method, along with the Seventh Sense™ biosampling device^18–21^. These devices collect larger volumes (>100 µL) than traditional finger-prick DBS sampling, and the sample is kept in liquid form rather than dried on paper, which is ideal for analytics that require a higher volume liquid sample, such as transcriptomic analyses. Moreover, these devices are simple to use, and users report less pain while using these compared to a finger prick or traditional venipuncture.^18, 19^ However, lancet-based blood collection devices, such as the Tasso-SST™, are more costly than a DBS sampling set up, and require further stabilization of liquid blood for certain analytes such as RNA.

There is an obvious advantage to decentralizing blood collection beyond a phlebotomy clinic; consequently, there have been many technological advances in self-blood collection. However, blood collection is often only the first step in blood-based laboratory tests. In a traditional outpatient or research setting, after a phlebotomist draws blood, it is processed soon after in a lab by a technician. Blood is not a static tissue sample; it contains living cells that can continue reacting to changes in their *ex vivo* environment. Post-collection sample handling is critical, and there have been numerous studies determining the best way to store, stabilize and handle blood samples for various target analytes.^22–24^ For transcriptomics profiling, RNA stabilization in liquid whole blood is particularly critical. Degradation of RNA by ribonucleases and rapid fluxes (induction and decay) of mRNA transcripts in response to the post-collection *ex vivo* environment can be highly unfavorable for research intended to understand *in vivo* cellular expression landscapes. Further, these *ex vivo* changes can lead to an inaccurate representation of the *in vivo* transcriptome in question^25, 26^

In a traditional outpatient venipuncture setting, stabilization of whole blood RNA is accomplished by collecting venous blood directly into vacutainers containing RNA stabilizers (e.g., Tempus™ or Paxgene™) or immediately pipetting anti-coagulated blood into RNA*later*™ containing vials. This procedure is incompatible with a self-sampling regime, as users cannot be expected to pipette their own blood or do venipuncture into a vacutainer tube on themselves. To fully enable remote sampling and transcriptomics profiling of liquid blood samples, one must eliminate the need for a phlebotomist-assisted blood draw *and* enable the patient or research participants to act as their own laboratory technician, allowing them to perform necessary steps to stabilize their blood sample without the use of pipettes, gloves, or syringes. In the present manuscript, we accomplish this goal of both collection *and* RNA stabilization. We combine a commercially available lancet-based blood sampling device (Tasso-SST™), a liquid RNA stabilizer (RNA*later*™), and a custom-engineered fluid transfer and stabilizer tube into a single sampling unit that can be mailed to study participants. Participants collect a liquid sample (∼ 0.1 - 0.5 mL) of whole blood, stabilize it, and ship it back to the laboratory for analysis. To demonstrate our liquid stabilization technology, we chose to target total whole blood cellular RNA with RNA*later*™. However, our technology is broadly generalizable in that a researcher interested in using a different stabilizer or targeting a different class of biomarkers can replace the stabilizer in the stabilizer tube with another liquid stabilizer.

To assess the usability and feasibility of this sampling methodology, we conducted a pilot study (*n =* 47 participants) to answer two fundamental research questions: 1) is the design and instructions for the kit comprehensive and user-friendly enough to allow users to collect and stabilize a sample of their own blood without in-person training?, and 2) is the stabilization process sufficient to enable isolation of high-quality RNA suitable for standard gene expression analyses? In the pilot study, the *home*RNA kit was sent to 47 participants, aged between 21 and 69 and living across 10 different US states (WA, CA, CO, NE, WI, IN, PA, NY, MA, ME). We demonstrated successful blood collection and RNA stabilization measured by total RNA yield and RNA integrity number (RIN). Additionally, we demonstrated expression analysis of two reference genes. Our kit and methodology open the potential for a new class of transcriptomics studies, enabling increased sampling frequency for longitudinal studies and access to populations that have been historically hard to reach.

## MATERIALS AND METHODS

Please see the supplemental information for the complete materials and methods, with information included on device fabrication, kit assembly, RNA analysis and participant demographics.

## RESULTS AND DISCUSSION

### Self-sampling kit for peripheral blood collection and RNA stabilization: overarching considerations of the *home*RNA blood kit

We developed *home*RNA to enable the self-collection of blood and immediate stabilization of RNA, an ability that opens new opportunities to probe immune responses to time- and location-specific stimuli outside of traditional venipuncture collection limitations. The two main components of *home*RNA include the Tasso-SST™, which collects approximately 100-500 μL of blood from the upper arm and a stabilizer tube, designed to screw onto the detachable blood collection tube from the Tasso- SST™ device. Figure 1A summarizes the general workflow for collection and stabilization using *home*RNA. To operate the Tasso-SST™, the device is first applied to the upper arm (Fig. 1A), where it is held in place by an adhesive. The user then presses a red activator button deploying a lancet that quickly punctures the skin. Blood is then drawn into a detachable collection tube, which holds up to approximately 500 μL of blood. To stabilize the freshly drawn blood, the collection tube containing the blood is detached from the Tasso-SST™ device and screwed tightly onto the stabilizer tube, and the connected tubes are shaken to thoroughly mix the blood with the stabilizer. The stabilized blood sample is then packaged and mailed back to the lab for analysis.

**Figure 1.**
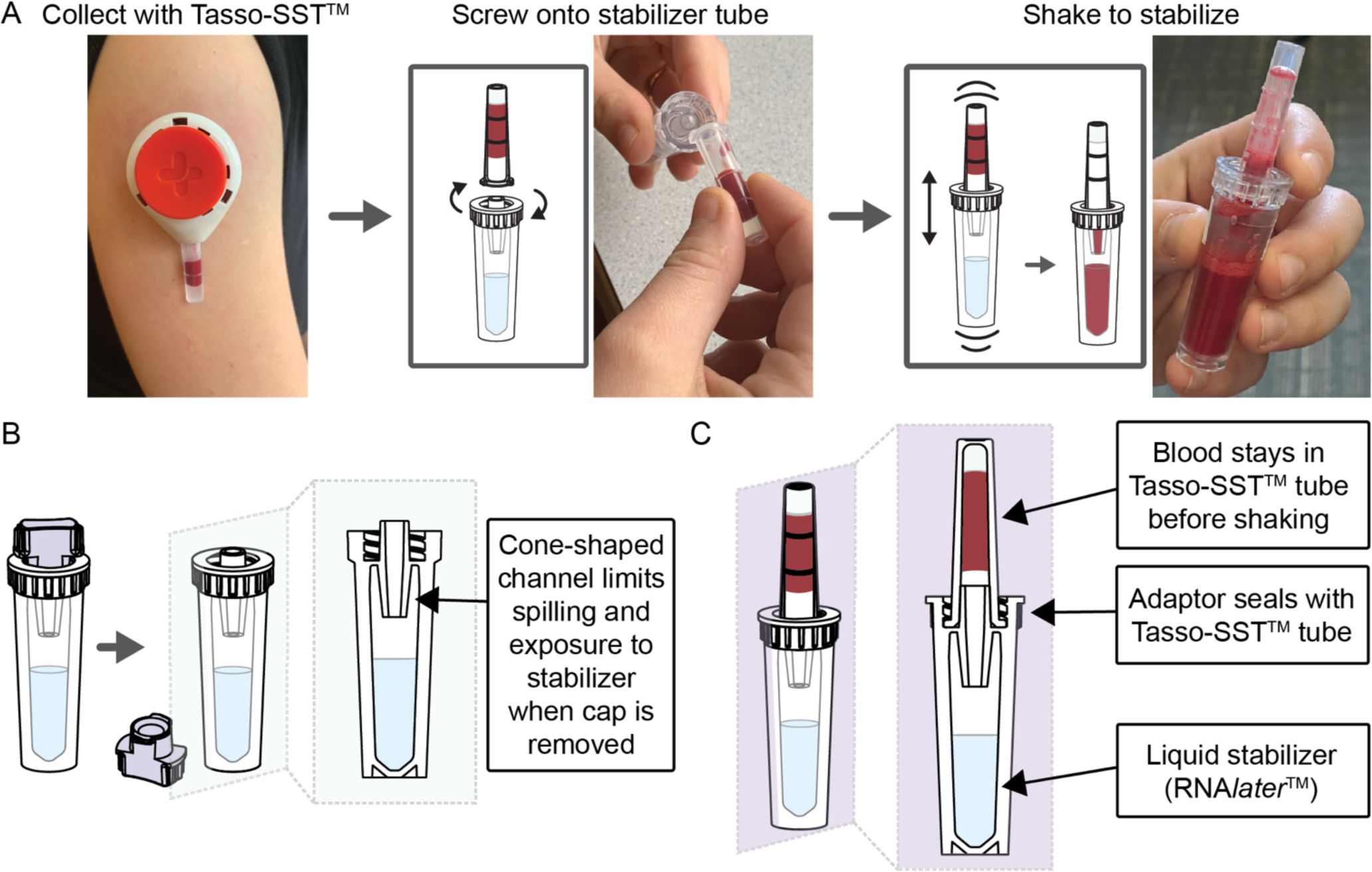
Workflow and design of *home*RNA blood collection and RNA stabilization kits. A) Workflow for *home*RNA, where blood is first collected using the Tasso-SST^TM^, screwed on tightly to the stabilizer tube, and then shaken to mix with the stabilizer, RNA*later*^TM^. B) Cross-sectional schematic of the stabilizer tube after the cap has been removed. C) Cross-sectional schematic depicting a Tasso-SST^TM^ blood collection tube filled with blood attached to the stabilizer tube, prior to shaking to mix the blood with the stabilizer.

The Tasso-SST™ was chosen as a method for blood collection due to its ability to be self-administered, its general ease of use, and the larger blood volume (>100μL) it draws when compared to other devices in its category, making it suitable for applications requiring a greater amount of starting material (e.g., RNA sequencing (RNA-Seq)). Compared to other blood collection methods, users report lower pain thresholds when using the Tasso- SST™ or similar devices that collect from the upper arm.^18–21^ The authors note that the serum separator tube (SST) gel (included in the Tasso-SST™ collection tube) is not necessary for RNA stabilization and analysis. In fact, a device or tube containing EDTA or another anticoagulant would be preferable to prevent clotting, and we note varying degrees of clotting observed in our returned samples. At the time of the study, the Tasso-SST™ was available for purchase. Therefore, it was chosen as the device to use as an initial proof of concept for demonstrating remote blood collection and RNA stabilization.

### Design of the stabilizer tube

The two primary design considerations for the stabilizer tube were 1) preventing exposure of the stabilizer solution and 2) preventing leaking or splashing when the blood and stabilizer solution are mixed. These design parameters are critical for home use, particularly to ensure that both the user’s blood and stabilizer solution remain contained within the tubes throughout the stabilization process. To address these design problems, we designed 1) a spill resistant stabilizer tube opening to mitigate user exposure (Fig. 1B), and 2) an opening that seals tightly with the Tasso-SST™ (Fig. 1C) to prevent leaking or splashing while mixing. Because of the small volume and shape (i.e., long and skinny) of the Tasso-SST™ blood collection tube, the blood sample remains in the tube even when it is inverted due to surface tension. This observation combined with a no-spill cone feature in the stabilizer tube allowed the user to easily tip the tubes sideways to connect them (Fig. 1A) without causing either liquid (the stabilizer or the blood sample) to spill. However, the blood’s tendency to remain in the Tasso-SST™ blood collection tube presented a non-trivial engineering challenge for mixing, since to be stabilized, the blood needed to interface with the stabilizer liquid in the opposite tube (Fig. 1C). However, when attached and shaken up and down, the surface tension is broken as the two liquid interfaces (the blood and the stabilizer) come into contact, allowing for mixing. Because this mechanism for mixing required vigorous shaking across the two tubes’ attachment point, a tight seal was critical. Leaking could also happen during the sample’s return if the seal was inadequate. To achieve a tight seal, the stabilizer tube piece that attaches to the Tasso-SST™ collection tube was based on the design of the cap included with the Tasso-SST™.

Surface tension was also utilized to design the opening of the stabilizer tube to achieve the first objective of preventing exposure to the stabilizer. The stabilizer tube was engineered with a fluidic cone-shaped channel at the connection point between the stabilizer tube and the Tasso-SST™ tube (Fig. 1B). This fluidic cone takes advantage of surface tension to create a valve such that the stabilizer solution remains in the stabilizer tube when the tube is inverted. Figure 1 illustrates schematic cross-sections of the stabilizer tube open (Fig. 1B) and attached to the Tasso-SST™ collection tube (Fig. 1C). The design of the stabilizer tube device was updated to facilitate easier mixing by altering the width of the opening at the bottom of the cone channel feature (Fig. S4). The wider opening also enabled easier pipetting of the blood sample from the tube during processing. A new glue was also used to allow for easier fabrication. The new design and glue were used in the last two groups of participants (groups 6 and 7). Additional details on the design and fabrication of the stabilizer tube can be found in the supplementary information.

All components that comprised the stabilizer tube (tube, adapter, and cap) were injection molded out of the same material (polycarbonate), to account for consistent material shrinkage during the injection molding process. Polycarbonate is commonly used in biological laboratory consumables and is known to be inert to most biological samples and reagents. Due to the nature of the injection molding process required for the adaptor piece (which included an internal thread feature), polystyrene and polypropylene, other commonly used materials in laboratory consumables, could not be used.

### Stabilization of blood using RNA*later*™ resulted in higher RNA yields and quality compared to other stabilizers over broad storage conditions

We assessed three RNA stabilizers commonly used in gene expression studies (Tempus^TM^, PAXgene^®^, or RNA*later*^TM^) for both yield and quality of the total RNA isolated from stabilized blood samples over broad storage conditions. Both Tempus^TM^ and PAXgene^®^ are often used in blood gene expression studies due to the commercial availability of these stabilizers in vacutainer tubes allowing for a direct draw of venous blood into the stabilizers. On the contrary, RNA*later*^TM^ is widely used to stabilize transcripts in laboratory specimens and extracted tissues but not commonly used in blood gene expression studies due to the lack of commercially available RNA*later*^TM^ vacutainer tubes. However, since any RNA stabilizer can be put in the stabilizer tube in the *home*RNA kits, we were not limited to vacutainer tubes and therefore assessed all three common stabilizers for preserving transcripts from blood. We assessed these three stabilizers for both storage temperature and length, the two major variables that could significantly affect post-collection RNA yield and quality. In this study, we measured RNA quality with an RNA integrity number (RIN) obtained on a Bioanalyzer 2100 coupled with its corresponding electropherogram profile. The RIN value is determined based on an algorithm that analyzes the electrophoretogram obtained from the capillary electrophoresis on the bioanalyzer chip. RIN values range from 1-10, where 10 represents entirely intact and non-degraded RNA.^27^ Annotated examples of an electrophoretogram and digital gel obtained from one of the samples from this study are provided in the SI in Figure S18. More information on the bioanalyzer, including examples of digital gels and electrophoretograms from samples with various RIN values can be found in Schroeder *et al* 2006.^27^ This part of the study was performed in-lab using blood collected from venous draws so that we could expose blood samples collected from one donor to controlled temperatures for fixed periods.

Preliminary experiments showed that blood stabilized in RNA*later*™ offered comparable total RNA yield and highest RIN values (yield = 4.9 µg RIN = 8.4) compared to both Tempus^TM^ (yield = 5.1 µg RIN = 7.1) and PAXgene^®^ (yield not detected, RIN = 1.0) after 7 days of storage at ambient temperature (Fig. S5). Given these preliminary results, both RNA*later*™ and Tempus^TM^ were assessed further for performance at a broader range of storage temperatures. As depicted in Fig. 2 (see Fig. S6 for electropherogram profiles), stabilization of blood using RNA*later*™ yielded better RIN values at higher temperatures. These parameters could be experienced with remote user-administered sampling methodologies. Furthermore, the lack of corrosive (tartaric acid) and toxic (guanidine hydrochloride) stabilizing chemicals in RNA*later*™ makes it an attractive choice for home-use or user- administered procedures. Due to high observed efficacy in stabilization for variable time and temperature profiles in our initial in-lab testing, coupled with user safety considerations, we chose to incorporate RNA*later*™ into the *home*RNA stabilizer tubes to accomplish stabilization of peripheral blood drawn with the Tasso-SST^TM^ in the pilot study.

**Figure 2.**
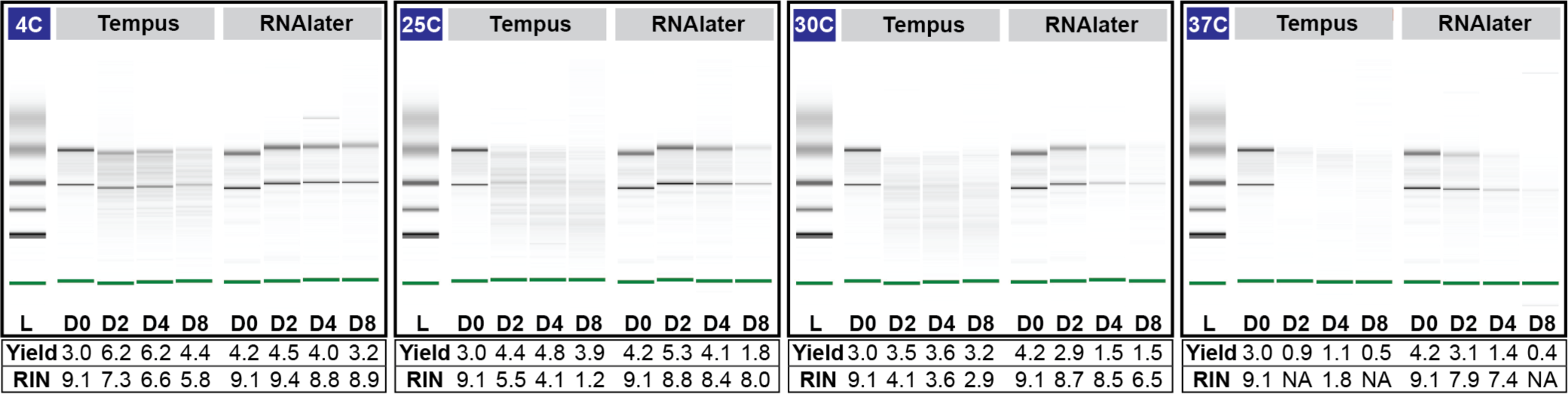
Performance of Tempus™ vs. RNA*later*™ on total RNA yield and RNA quality. Effect of storage temperature and duration on total RNA yield (μg) and quality (RIN value) of samples stabilized in Tempus™ vs. RNA*later*™. Digital gel image of Tempus and RNA*later*™ stabilized blood stored at 4°C, 25°C, 30°C and 37°C over 8 days.

### Analysis of whole-blood RNA returned from the *home*RNA kit reveals feasibility for disseminated whole blood sampling and RNA stabilization

We enrolled 47 participants in a pilot feasibility and usability study to demonstrate self-blood collection and RNA stabilization outside of a clinical or research setting. This study was approved by the University of Washington Institutional Review Board (IRB) under protocol STUDY00007868. All study procedures were performed after informed consent was obtained. Participants were asked to use either one or two *home*RNA kits each time they were sent kits, and some participants were sent kits on more than one occasion. With these considerations, 47 participants generated 60 samples. Details of which participants used one or two kits is detailed in the SI (Table S3), and further details of which participants were included in two groups and how many samples came from each participant are included in the Supplemental Dataset. *home*RNA kits were mailed to the participant’s home where they collected, stabilized, and returned their own blood sample based solely on provided instructions in the kits and instructional video (see SI). Stabilized samples were returned to the laboratory for analysis via mail. Therefore, they had to remain stable throughout the shipping and the variability of temperature, pressure changes, and other mechanical stress inflicted during shipping.

Upon returning to the lab, total RNA was extracted from all stabilized blood samples and assessed for yield and quality (RIN values). 83% and 100% (*n* = 60 total samples) of blood samples returned from the pilot study offered a total RNA yield greater than 500 ng (a comfortable minimal cut-off value for large-scale transcriptomics analyses) and 100 ng (a comfortable minimal cut-off value for expression analyses of a small panel of targeted genes), respectively (Fig. 3A). These cut-off values for total yield obtained immediately after extraction may vary across studies depending on the choice of analysis methods (e.g., RT-PCR, digital droplet PCR, RNA-Seq, xMAP^®^ and nCounter^®^ technologies) and pre-analysis sample processing steps (e.g., globin depletion, RNA species enrichments) that will incur further yield losses. Based on our pilot study data, all self-drawn and self-stabilized peripheral blood samples using the *home*RNA blood kit offered sufficient yield for targeted small gene panel profiling. The majority of the samples (83%, *n* = 50/60) also have sufficient yield for genome-wide transcriptional profiling analysis methods such as RNA-Seq.

**Figure 3.**
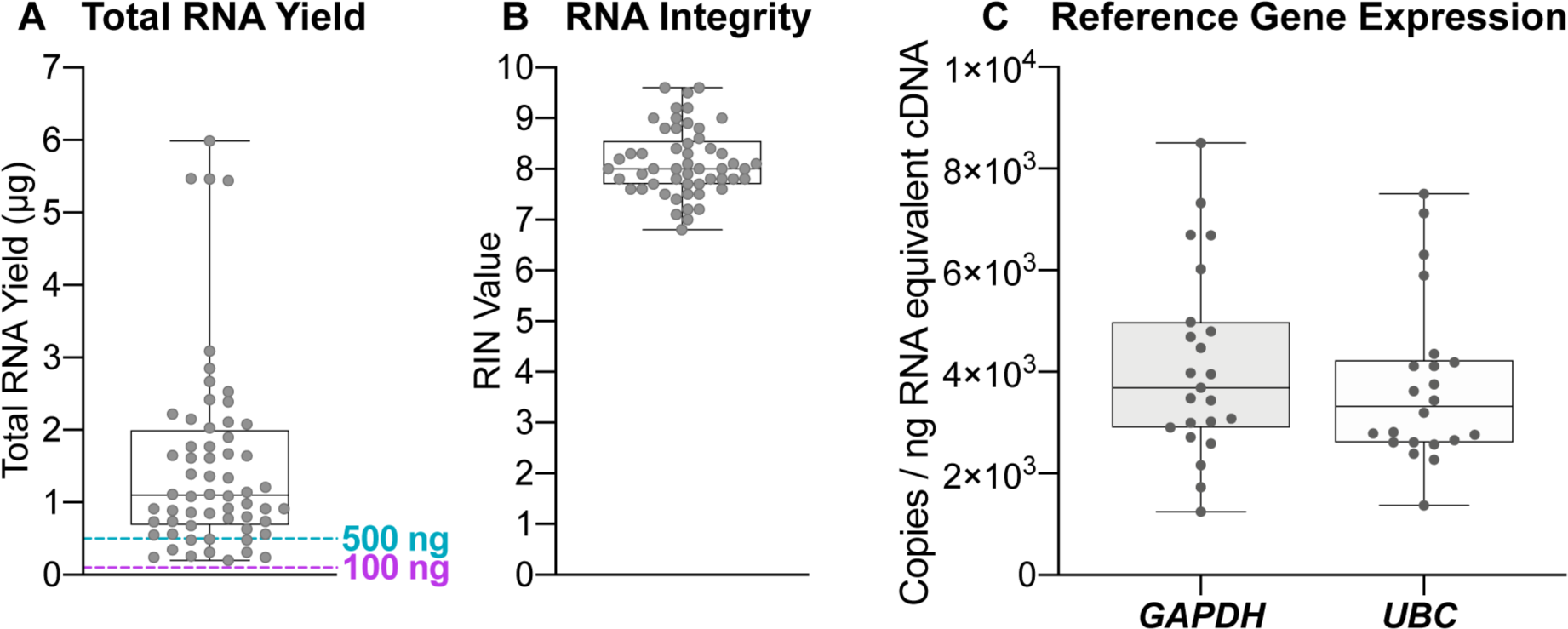
RNA yield and RIN value of peripheral blood samples self-collected and stabilized with *home*RNA. A) Total RNA yield (μg) and B) RIN value and C) droplet digital PCR of *GAPDH* and *UBC* from RNA isolated from peripheral blood samples collected and stabilized by participants in their home using *home*RNA.

We obtained RIN values for 85% of all isolated RNA (*n* = 51/60) samples (Fig. 3B). We note that the 15% (*n =* 9/60) of samples that were not scorable and did not afford a RIN value primarily due to the low total yield, resulting in low RNA concentrations in these samples (Fig. S16). For samples that did not afford RIN values, a visual inspection of the digital gel images of these samples showed 78% (*n =* 7/9) contain intact ribosomal RNA bands depicting good RNA integrity (Fig. S17). RIN values of all scorable RNA samples range between 6.8 – 9.6, with 53% (*n* = 27/51) of samples affording RIN values greater than or equal to 8.0 and all but one of the samples greater than 7.0 (*n* = 50/51) (Fig. 3B). Similar to the minimum cut-off value for yield, RIN values and their suitability for downstream gene expression analyses vary widely based on the source of the tissue or sample from where the RNA is isolated. For example, formalin-fixed paraformaldehyde embedded (FFPE) tissues and tissues containing high levels of ribonucleases (e.g., blood, liver, spleen, and kidney tissues) often afford lower RIN values due to the high degradation potential in these tissue types. A RIN value of 7.0 is a typical minimal cut-off value for RNA-Seq applications.^17, 28^ For highly degraded samples such as FFPE tissues that often have RIN values as low as 2.0, the fragment size distribution index (DV200) is frequently used to assess RNA quality and determine sample suitability for downstream gene expression analyses.^29–30^ Further, as there is interest in using RNA-Seq for lower yield or degraded samples (such as FFPE), there are many published techniques on methods to accomplish RNA-Seq in degraded samples.^28, 31^ Despite whole blood being rich in ribonucleases, the remote self-collection and stabilization process of the *home*RNA blood kit still afforded high RIN values (RIN > 7.0) that, by themselves, render these samples suitable for a variety of gene expression analysis. Therefore, a DV200 index assessment was not necessary for assessment of these samples. The yield and quality of the RNA extracted from blood samples from our pilot study are well within the parameters for targeted small gene panel profiling, and many of the samples even reach the higher thresholds set by sequencing facilities, conferring the convenience of outsourcing RNA-Seq for researchers interested in using our *home*RNA kit for their own out-of-clinic transcriptomics studies. Importantly, when compared to emerging remote self-sampling methodologies such as dried blood spotting, the *home*RNA collection process affords a better yield and quality that allows for a broader range of flexibility in analysis methodologies.^17, 32, 33^

Despite sufficient yield and high RIN values observed for the RNA samples isolated from the *home*RNA stabilized blood, presence of other residual chemical impurities that may have been introduced during the assembly process (e.g., the epoxy used for bonding) may affect downstream gene expression analyses. Thus, to further assess whether RNA isolated from *home*RNA-blood samples are compatible with downstream gene expression analyses protocols, expression of two reference genes (*GAPDH* and *UBC*) were measured from isolated RNA samples (*n* = 23) using digital droplet PCR (ddPCR). ddPCR was chosen as an analysis method due to its high protocol similarity to commonly used RT-PCR methods coupled with additional protocol requirements of maintaining droplet stability upon droplet generation. Select RNA samples with yield (0.63 – 5.44 µg) and RIN values (7.1 – 9.6) used for ddPCR analysis broadly represent the range of values for each of the two parameters observed within our pilot study. As shown in Fig. S7, the mean values of total accepted droplets (TAD) for both *GAPDH* (TAD = 16,203; *n* = 23) and *UBC* (TAD = 16,693; *n* = 22) reactions are comparable to that of the no-template control (NTC) (TAD = 16,741; *n* = 4), suggesting droplet stability is maintained throughout the amplification process. Both *GAPDH* and *UBC* depicted mean (SD) values of 4.1 x 10^3^ (1.9 x 10^3^) and 3.7 x 10^3^ (1.6 x 10^3^) copies/ng RNA-equivalent cDNA respectively (Fig. 3C). The variation in observed copies can be attributed to biological variations in *GAPDH* and *UBC* gene expression within the study participants. Taken together, RNA yield, quality, and gene expression results obtained from *home*RNA blood RNA samples demonstrated successful preservation and expression analysis of blood mRNA transcripts from this user-operated home sampling methodology. Future studies on the use of the *home*RNA decentralized blood collection and stabilization technology to capture dynamic changes of immune responses to a variety of environmental stimuli and diseases will be of utmost interest to our group and the broader community.

### Kit performance is robust across participant demography and mailing groups

To assess the geographic distribution feasibility of this remote-sampling methodology, *home*RNA blood kits were mailed from our lab (Seattle, WA) to various residential destinations in the West Coast, Midwest, and East Coast of the United States (Fig. S8). Additionally, we mailed *home*RNA to various residential housing types in urban, suburban, and rural areas, including single-family unit houses and large multi-family apartment complexes (where packages are typically held in the lobby or mailroom). Demonstrating sampling from rural areas is important to expand research in places where participants traditionally needed to travel to phlebotomy labs located in other towns or cities. Such expansion would enable research into immune events that may be more commonly triggered in rural populations, such as exposure to agricultural chemicals or wildfires.

*home*RNA kits were sent out in seven independent mailing groups from May to December 2020 (Table S3). Most kits were returned within 1-3 days after sample collection, but some kits were returned later, with one being returned as late as 15 days after collection (see supplemental dataset). Notably, this sample still yielded an RNA yield of 0.56 µg and a RIN value of 7.8, so even with over a two-week delay, where the sample was in the mail, we recovered enough intact RNA for downstream analyses. From a usability perspective, differences between groups in terms of total RNA yield and RIN values were minimal (Fig. S9), suggesting robustness to the remote sampling methodology afforded by the *home*RNA kit; slight variations in the instructions (e.g., changes in wording, updated graphics, the inclusion of an instructional video) did not dramatically change results, suggesting the kit itself was relatively simple and intuitive to use. Finally, the RNA quality analysis parameters (total RNA yield and RIN values) were not markedly different across a range of age group (Fig. S10), gender (Fig. S11), or body mass index (BMI) (Fig. S12). Reported blood levels (which we use as an approximation for volume collected) also were not markedly different between different ages, genders, or BMI (Fig. S13), indicating consistency in blood collection with the Tasso SST™ regardless of participant demographic or BMI. While we demonstrated robustness across these parameters, we note that the pilot study did not collect information on participants’ socioeconomic status or level of education. For future studies, we intend to evaluate the usability of the *home*RNA kit across a more diverse population and in geographical regions or during seasons that can incur more considerable variations in the high/low ambient temperatures, as a notable limitation in our geographical sample set is that kits were not mailed to the hotter regions of the United States (e.g., the Southwest or South) during the summer months, and the majority of the samples were taken from the Seattle region; we are addressing this limitation in our ongoing studies.

### Participant survey responses indicate good usability of *home*RNA

Ease of use and participant perception is critical for compliance, particularly for using this method in future longitudinal (multi-sample) studies. Usability was assessed through a user experience survey that the participants were asked to complete after using the kits. This survey was also used as a mechanism for feedback from the participants in order to iterate upon the instructions and kit components. Perception of the kits in terms of the time it takes to complete it, ease of use, and pain or discomfort were assessed (Fig. 4). Metrics for measuring the performance of the kit were also assessed, including asking the participants to estimate how much blood was collected and the time the Tasso-SST™ was left on the arm, as an approximation for the blood collection time (Fig. 5).

**Figure 4.**
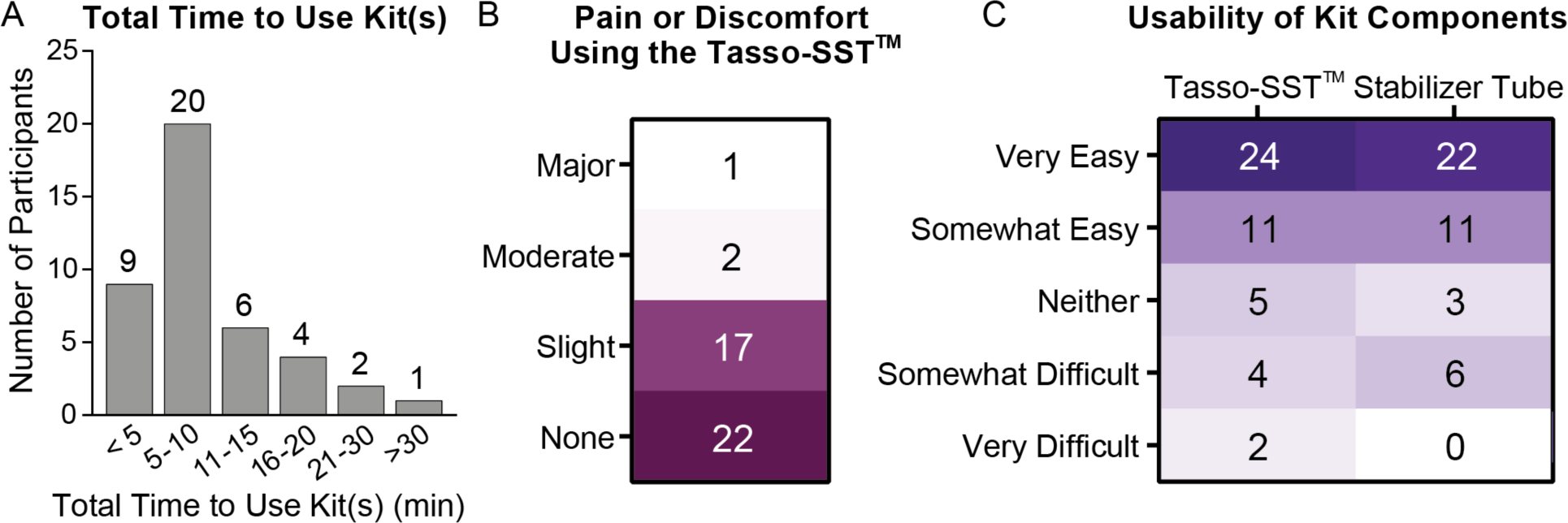
Survey responses to assess kit usability. A) Histogram showing total self-estimated time required for each participant to use the kit from start to finish. B) Participant ratings for level of pain or discomfort using the Tasso-SST™. Numbers represent number of participants. C) Participant ratings for the usability of the different kit components. Numbers represent number of participants.

**Figure 5.**
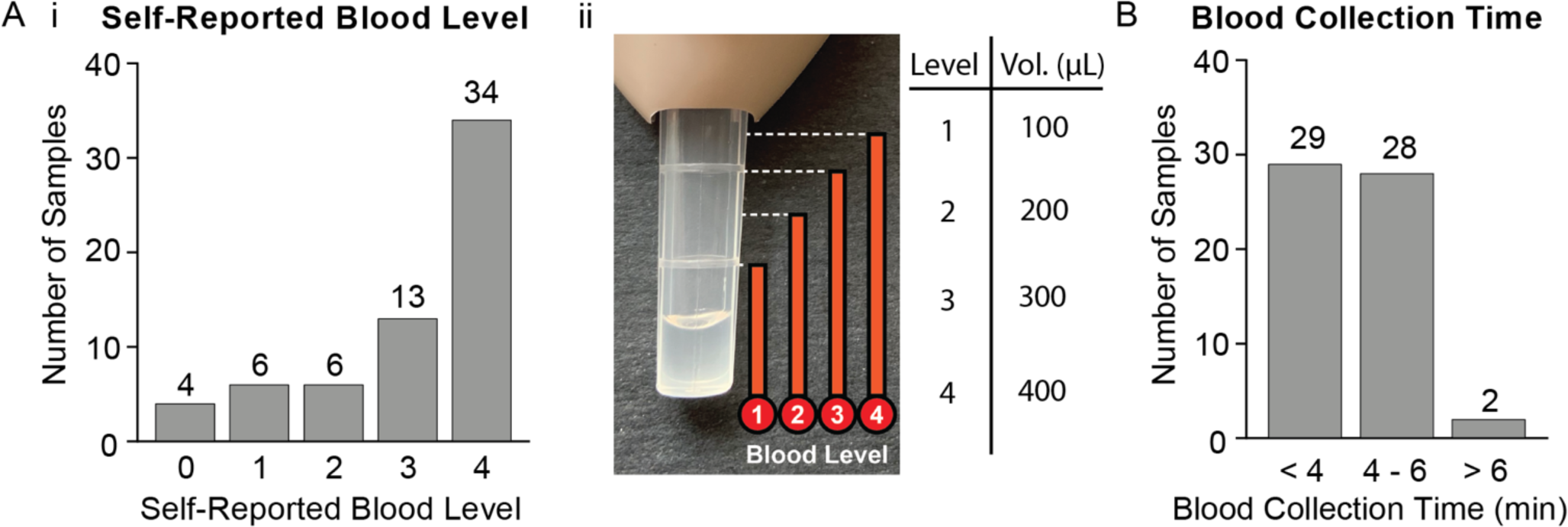
Survey responses to assess kit performance. A) i) Participant responses when asked to estimate the approximate blood level they filled the Tasso- SST™ blood collection tube based on the ii) picture provided in the online survey. The volumes corresponding to each level are noted in Aii. B) Participant responses when asked how long they left the Tasso-SST™ blood collection tube on their arm - this includes time before collection and after collection has stopped before removal of the device.

Most of the participants who successfully collected blood using the Tasso-SST™ finished using their kits in less than 10 minutes (69%, *n =* 29/42, Fig. 4A). Nearly all participants (93%, *n =* 39/42) reported either minimal pain or discomfort or no pain or discomfort while using the Tasso-SST™, and the majority (52%, *n* = 22/42) reported no pain or discomfort. Only 3 participants found the pain or discomfort to be rated as moderate (*n* = 2) or major (*n* = 1). Regarding the participant’s perception of how easy the kits were to use, most of the participants reported the Tasso-SST™ and stabilizer tube to be easy to use or somewhat easy to use (76%, *n* = 35/46 and 79%, *n* = 36/42 for the Tasso-SST™ and stabilizer tube respectively). Participants with close proximity to the project were excluded from this analysis; participants who failed to collect blood were excluded from the pain and stabilizer tube usability but were included in the usability for the Tasso-SST™ (4 out of 47 total enrollment). In summary, feedback with regards to the pain experienced and usability was positive.

### Participant survey responses on device performance show sufficient yield even at a low sample volume

To assess the possible correlation between estimated blood collection volume and total RNA yield, participants were asked to estimate the levels of blood drawn into the Tasso-SST^TM^ blood collection tube based on a provided blood tube image (depicted in Fig. 5Aii). Estimated volumes for the four levels are as follow: Level 1 = 100 μL, Level 2 = 200 μL, Level 3 = 300 μL and Level 4 > 400 μL. The majority of the participants reported blood collection at Level 4 (52% *n* = 34/65). The reported blood level versus the RNA yield is plotted in Fig. S14A. While there are very few samples at lower reported blood levels (Level 1 and 2), there does not appear to be a strong correlation between blood level reported and yield. This could be due to inaccuracies in reporting by participants, individual variability in RNA yield, or loss during the sample processing or blood clotting in the stabilizer tube. Notably, participants who reported collecting a low volume (Level 1 ∼100 μL) of blood still had RNA yields >100 ng, with one sample as high as 2.85 μg. We have also included data on blood collection time, reported blood volume, and RNA yield in the SI (Fig. S14B and S15). In short, all collections, irrespective of reported collection levels, afforded RNA yield sufficient for downstream gene expression analyses.

## CONCLUSION

Our *home*RNA kit will enable translational researchers to ask fundamentally different biological and clinical questions than have to-date been limited to clinic-based transcriptomics. Given the flexibility of this sampling system and low sample volume compared to venipuncture, future studies involving frequent sampling and sampling around a specific event (e.g., disease flare, environmental or pathogen exposure) can be employed to better elucidate hard-to-capture expression signatures of the immune response. For example, we hope that our sampling platform enables us to observe and study early, transient, and dynamic changes in both the innate and adaptive arm of the immune system throughout the various stages of an infection in order to guide treatment or transmission control measures.

Critically, with *home*RNA, multiple samples can be taken from the same individual outside of an in-patient setting more readily than with current methods, enabling easier comparison to an individual’s own baseline. Coupling this with the ability to sample virtually anywhere, studies into a person’s individualized response (that is, compared to their own baseline) to an exposure or event in their daily environment are possible. Finally, we are excited to expand this technology to disseminated diagnostics, therapeutics, and clinical research into lower resource or rural settings, which are often far from a phlebotomy clinic.

## ACKNOWLEDGEMENTS

This publication was supported by the David and Lucile Packard Foundation (for device manufacturing and human subjects components of the research); the University of Washington; and the National Institutes of Health (NIH) through National Institute of General Medical Sciences award number R35GM128648 (for the in-lab developments) and by the National Center For Advancing Translational Sciences award number UL1 TR002319. The content is solely the responsibility of the authors and does not necessarily represent the official views of the National Institutes of Health. We would like to acknowledge Ulri Lee for helpful discussions. We would also like to thank Lochlan Hickok and Paul Miller for help facilitating shipping and logistics.

## *home*RNA KIT AVAILABILITY

For the purposes of repeating this work, the authors will endeavor to make the *home*RNA kit available to other research groups at production cost, provided supplies and resources allow. Please contact the corresponding author for more information.

