## Supplemental Information for "*home*RNA: A self-sampling kit for the collection of peripheral blood and stabilization of RNA"

### Table of Contents:

Materials and Methods Pg. S2 – S6
Supplemental Tables and Figures Pg. S6 – S19
References Pg. S19
Instructions for Use Pg. S20 – S21

**Supplementary Information not included in this document:**

Supplemental Dataset

Contact corresponding author for design files for tube components (Adaptor 1, Adaptor 2, Cap, Vial) and sample holder:

#### MATERIALS AND METHODS:

##### Development and assembly of the homeRNA blood collection and stabilization kit

*Design process for the RNA stabilizer tube:*

The RNA stabilizer tube is assembled from three individual components: 1) a reagent vial to house the liquid stabilizer 2) an adaptor that interfaces both the reagent vial and the Tasso-SST™ blood tube and 3) a vial closure cap. Preliminary design iterations were generated on SolidWorks and 3D printed on a Form 3 3D printer (Formlabs) using clear resin (Formlabs). These preliminary designs were evaluated for leakage and spillage when tipped over or agitated. The mechanical agitation of shipping was simulated by placing the tube designs filled with water in a box and shaking. Designs were also tipped over without caps on to evaluate if fluid would easily spill out of the tube if an end user were to accidentally tip them over. The tubes were then inspected for leaking or spilling. The cone-shaped channel design, which was ultimately used, performed notably better on leaking and spilling metrics in both the simulated shipping and knocking over tests, compared to a straight channel shape, or an inverted cone shaped channel where the opening on the bottom was wider than the top. The design of the adaptor piece was altered partway through the study (between group 5 and 6) to allow for easier mixing by increasing the diameter of the cone feature (Fig. S4). The increased diameter also allowed for easier pipetting of the returned blood sample; the blood sample could be safely pipetted using a polypropylene Pasteur pipette (CELLTREAT) with a 2 mL manually adjustable pipette (Socorex) and a Pasteur pipette adaptor (DWK Life Sciences). Specific features of the stabilizer tube design include a cone channel feature in the adaptor piece (Fig. 1B) to prevent contact between stabilizer reagent and the user during blood stabilization.

*Design considerations for injection molding of the RNA stabilizer tube:*

Both tube designs were injection molded out of polycarbonate (PC: Makrolon 2407) by Protolabs, Inc (Maple Plain, MN). Several design considerations were optimized to include features important for the process of injection molding. Briefly these include, 1) adding a 2 degree draft to all vertical surfaces on all parts to allow for easier removal of the part from the mold, 2) thickening of the adaptor and vial piece to allow for mold ejection pin zones, 3) designing the shape of the cap piece to avoid overhangs, allowing for a two piece mold, and 4) designing the internal threads on adaptor piece, such that it required only a quarter turn to remove from the mold. Polycarbonate was chosen as the material for all three pieces as a consideration for constant material shrinkage, as the parts needed to fit together. Polycarbonate was chosen over other commonly used plastics due to the constraint on material caused by the internal thread feature in the adaptor piece; we note that polycarbonate is also a common material for biochemical samples containers.

*Fabrication of the RNA stabilizer tube:*

For assembly, all components of the stabilizer tube are first cleaned via sonication in 70% ethanol (v/v) for 30 min and air dried. The adaptor was bonded onto the reagent vial using a UV curing glue (Damn Good® for the first 5 groups and Dymax MD® 1450-M-UR-SC for groups 6 and 7, Fig. S3). The Dymax UV curing glue was ultimately chosen for the last iteration due to several design considerations. It has a lower viscosity making it suitable for fluid manipulation with a standard micropipette, enabling faster and more consistent manual application of the glue to the bonding line. Further, the Dymax glue contains a color change indictor to aid in visualization for curing. Bonded parts were cured for 60 min at

395-405 nm UV Lamp (Quans) for all tubes used in all groups. The stabilizer tube was filled with 1.4 mL of RNA*later*™ (Thermo Fisher) as the stabilizing reagent, capped, and checked for leakage due to bonding defects before distribution to study participants. We did not observe any bonding defect leakage in tubes prepared for this study. RNA stabilizer tubes used in the feasibility pilot study were prepared within a week of being mailed to study participants.

*Evaporative loss from stabilizer tubes:*

To assess evaporative loss, 8 stabilizer tubes were filled with 1.4 mL of RNA*later*™ and capped and left at room temperature for 8 weeks. Tubes were massed during initial setup before and after addition of RNA*later*™*,* and were subsequently massed at week 1, 2, 3, 4, 5, and 8 (Fig. S19). Less than 1% loss of RNA RNA*later*™ mass occurred in all 8 tubes at the 1-week timepoint, and therefore a week was chosen as the cutoff as experiments on evaporation of RNA*later*™ from prepared and capped tubes indicated less than 1% evaporative loss (Fig. S19); in subsequent work, we have found that tubes filled with RNA*later* can be kept for 4 weeks while still allowing <5% v/v evaporative loss of RNA*later* (our 1-week cutoff in this study is unnecessarily stringent).

*Kit components and assembly:*

All components included in the *home*RNA blood kit are listed in Table S1 and shown in Fig. S1. The RNA stabilizer tube was labeled with a unique sample code and packaged into a transport bag with an absorbent material. A heat pack was included to increase blood flow to the upper arm (Medline). The stabilizer tube insert was designed to immobilize the stabilizer vial containing blood in a 50 mL conical tube during transport. The insert was designed on SolidWorks and 3D printed on a Form 3 3D printer (Formlabs) using clear resin. All kit components were placed in a rigid custom design mailer box fabricated via die-cutting (The BoxMaker, Inc.). A temperature strip (Propagate Pro) was affixed onto the outer side of the mailer box for temperature recording by study participants. A mailer bag with pre-printed return shipping label was also included with the outgoing package for specimen return to the lab. In brief, iterations were made to improve overall usability of the kit and clarity of the instructions for use. We included the final version of the instructions for use (IFU) at the end of this document.

*User experience survey:*

Subjects were asked to fill out a user experience survey after using their kit. The survey included questions about the timing in which they use the kit, the ambient temperature during use of the kit, the volume of blood collected based on how visually full the Tasso-SST™ blood tube was, and asked participants to rate their experience using the kit and provide any additional feedback on the kit or the process of receiving and returning the kit.

*Instructions for Use:*

The final version of the instructions for use (IFU) is found at the end of this document. The IFU version published here was not used with any of the 7 sample groups, but rather the most mature version that was developed based on user feedback from the pilot study. The IFU was also designed by the authors of this study based in part on images from www.tassoinc.com available at the time of our study. These IFU are for the *home*RNA kit only and are not vetted by Tasso Inc.

*Instructional video:*

Some groups (4-7) had the option to access an instructional video (linked to a QR code on their IFU). This is a 4 minute and 10 second video which features a member of our research group demonstrating proper use of the blood sampling and RNA stabilization kit. There is a voice-over narration describing each step in detail. The QR code in the posted IFU links to the video and a link to the video can also be found here: https://youtu.be/iV3GZ8SmmuM

**RNA stabilization, isolation, and gene expression analysis**

*RNA stabilization and isolation:*

*For in-lab RNA stabilization experiments*, fresh (< 8 hours post blood draw) whole venous blood drawn into EDTA-coated vacutainers (BD) was purchased from Bloodworks Northwest (Seattle, WA). Upon receipt, anti-coagulated whole blood was promptly transferred into either Tempus™, PAXgene®, or RNA*later*™ stabilizing reagents at their manufacturer's recommended stabilizer : blood ratios (v/v) of 2:1, 2:1, and 2.6:1 respectively. Stabilized blood was then incubated at various temperature ranges (4ºC, ambient, 30ºC, and 37ºC) and for a range of time periods (0-8 days) specific to each experiment as described in the results. At the end of the incubation period, total RNA was isolated from Tempus™-, PAXgene®-, and RNA*later*™- stabilized blood using Tempus™ Spin RNA Isolation Kit (Thermo Fisher), PAXgene® Blood RNA Kit (PreAnalytiX), and Ribopure™ - Blood RNA Isolation Kit (Thermo Fisher), respectively, according to manufacturer's protocol, and eluted in 50-100 µL volume.

*For the at-home blood collection and stabilization feasibility study*, Tasso-SST™ collected blood was stabilized in RNA*later*™ by the human subjects using the RNA stabilizer tube that interfaces with the Tasso-SST™ blood tube (Fig. S2). Total RNA was isolated using the Ribopure™ - Blood RNA Isolation Kit (Thermo Fisher) according to manufacturer's protocol and eluted in 50-100 µL volume. RNA concentrations were obtained on a NanoDrop® ND-1000 spectrophotometer (Thermo Scientific). RNA integrity number (RIN) values were obtained on a Bioanalyzer 2100 (Agilent) using the RNA 6000 Nano Kit (Agilent) following the manufacturer's protocol. Initially, 1 µL of isolated RNA was added to the bioanalyzer chip to obtain a RIN value as described by the manufacturer's protocol. If a RIN was not obtained the sample was run again with 2 µL of isolated RNA. Details as to which RIN values were obtained with 1 µL or 2 µL is located in the supplemental dataset file included in the supplemental material. Details on interpreting bioanalyzer data can be found in Schroeder *et al* 2006,[1] and a figure annotating the main components of an electrophoretogram profile and digital gel electrophoresis image of one of the samples collected in this study can be found in Fig. S18. Isolated RNA was stored at -80ºC until ready for further analyses.

*Digital droplet PCR analysis:*

1-3 µg of isolated RNA was digested with DNaseI (NEB) and 0.5 – 1.0 µg of DNA-free RNA was reverse transcribed into cDNA using Bio-Rad iScript™ cDNA Synthesis Kit (#1708891) according to manufacturer's protocol. ddPCR™ reactions were carried out in a total volume of 20 µL containing 10 µL of Bio-Rad 2X QX200™ ddPCR™ Evagreen ddPCR™ Supermix (#1864034), 25 ng of RNA-equivalent cDNA, and 180 nM of both forward and reverse primers (IDT). Primer sequences of both reference genes (*GAPDH* and *UBC*) analyzed in this study are listed in Table S4. Droplets were generated from the above reaction on a QX200™ droplet generator (Bio-Rad) using Bio-Rad QX200™ Droplet Generation Oil for Evagreen (#1864005) and Bio-Rad DG8™ cartridge (#1864008) and gasket (#1863009). 40 µL of the droplet suspensions were transferred to a Bio-Rad ddPCR™ 96-well semi-skirted plate (#12001925), sealed on a PX1™ (Bio-Rad) with a pierceable foil heat seal (#1814040) and cycled on a C1000 Touch™ 96-deep well reaction module thermal cycler (Bio-Rad). Thermal cycling conditions were as follows: initial denaturation and enzyme activation (95 ºC, 5 mins), 40 cycles of denaturation (95 ºC, 30 sec) followed by annealing and extension (55 ºC, 1 min) at a 2 ºC / sec ramp rate, signal stabilization (4 ºC, 5 min) and enzyme deactivation (90 ºC, 5 min), and finally held at 4 ºC indefinitely. After PCR amplification, cycled droplets were read promptly on a QX200™ Droplet Reader (Bio-Rad). Extraction of raw fluorescence amplitude data, quantification of positive/negative droplets, and droplet visualization were performed using Bio-Rad's QuantaSoft™ software.

**Pilot study: feasibility and usability assessment of the *home*RNA kit**

*Participant characteristics:*

This study was approved by the University of Washington Institutional Review Board (IRB) under protocol STUDY00007868. All study procedures were performed after informed consent was obtained. A total of 47 healthy volunteers between ages 21-69 years old were recruited via word of mouth or email to participate in the pilot study.

*Human subjects study design:*

**General study design:** Study participants were enrolled in groups in order to iterate on the general usability of the *home*RNA kit, specifically on the kit components and the clarity of the IFU (Table S2). The study enrolled a total of five groups. In each group, participants were asked to self-collect and stabilize blood from their upper arm using the *home*RNA blood collection and stabilization kit. Each participant was also asked to complete a user experience survey that was designed to guide further improvements to kit components and sampling parameters (e.g., ease of use, clarity of the IFU, mailing logistics, ambient temperature at collection site, etc.) Based on feedback, improvements were implemented in subsequent groups. Participants were asked to package their stabilized blood samples to be returned to the lab for analysis using the provided return mailer bag. Except for samples from group 1, where samples were picked up by the study team, all stabilized blood samples were mailed using next day delivery courier services (UPS). Unless specified, returned samples are stored at -20ºC until ready for RNA extraction. Study data were collected and managed using REDCap electronic data capture tools[2] hosted at the Institute of Translational Health Sciences. REDCap (Research Electronic Data Capture) is a secure, web-based application designed to support data capture for research studies, providing: 1) an intuitive interface for validated data entry; 2) audit trails for tracking data manipulation and export procedures; 3) automated export procedures for seamless data downloads to common statistical packages; and 4) procedures for importing data from external sources.

**Group-specific study design:** In order to assess effect(s) of storage length on total RNA yield and quality, participants in groups 1 ($n$ = 4) and 2 ($n$ = 5) and 7 ($n$ = 5) were given two *home*RNA kits to be used, one on each arm. Participants were instructed to use the second kit promptly upon completion of the first kit and to complete a user experience survey. For group one and two, one vial of stabilized blood from each participant was frozen at -20ºC immediately upon return to the lab, while the other was stored at ambient temperature for an additional three days prior to RNA extraction. Participants in groups 3-6 were given one *home*RNA kit and asked to perform the collection and stabilization procedure and complete a user experience survey. The first three groups were recruited locally (greater Seattle area) as well as group 6. To assess feasibility of the sampling pipeline across a wider geographical distribution, individuals from across the contiguous United States were recruited as participants for groups 4 ($n$ = 13), 5 ($n$ = 16) and 7 ($n$ = 5). Some participants were included in two groups. Specific information as to which participants were included in which groups as well as how many collected samples came from each individual can be found in the supplemental dataset. Taking into account participants completing two samples in one group and participant inclusion in two groups, 47 participants yielded 60 samples.

*Inclusion and exclusion criteria for subject enrollment:*

Inclusion criteria included aged 18-75 years, between 105 - 230 lbs for those assigned female at birth and 135 - 250 lbs for those assigned male at birth. Exclusion criteria included pregnant individuals or those currently breastfeeding, individuals with skin disorders such as scabbing or psoriasis located on the upper arm, individuals with a blood platelet or coagulation disorder, or currently taking any blood platelet or anticoagulant medications, individuals who are immunosuppressed or taking any immunosuppressive medication, or individuals who reside in a correctional facility.

*Recruitment and enrollment of subjects:*

Subjects were recruited by word of mouth, email, or posts on digital platforms advertising the study and invited to follow a link to a pre-screening survey which asked questions about the inclusion and exclusion criteria. A study team member then called the subject to confirm they met the inclusion and exclusion criteria and to answer any questions about the study. After signing an informed consent form on DocuSign, subjects enrolled into the study with an online form.

**SUPPLEMENTAL TABLES AND FIGURES:**

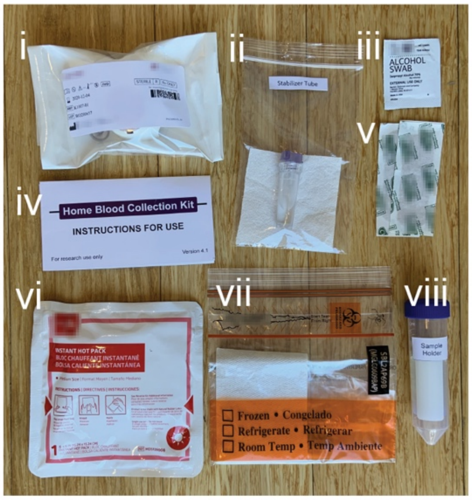


**Figure S1. Components of the *home*RNA kit.** Kit components include i) the Tasso-SST™ device, ii) the stabilizer tube containing RNA*later*™ iii) alcohol wipes iv) instructions for use, v) sterile bandages, vi) hot pack for warming the arm prior to application of the Tasso-SST™, vii) sample return bag and viii) sample holder with 3D printed insert to hold the sample tube in place.

**Table S1. Components of the *home*RNA blood kit.**

| Kit Component | Manufacturer(s) | Quantity |
|---|---|---|
| Sterile Tasso-SST™ blood collection device | Tasso, Inc. | 1 |
| RNA stabilizer tube | Our Lab | 1 |
| Instant heat pack | Medline Industries, Inc. | 1 |
| Sterile Alcohol Wipe | Covidien, BD | 2 |
| Sterile bandage | Band-Aid, Curad | 1 |
| Specimen transport bag with absorbent pad | Minigrip | 1 |
| 50 mL conical tube with stabilizer tube insert | BD, Our Lab | 1 |
| Instructions for use | Our Lab | 1 |
| Blood-stabilizer mixing instruction card | Our Lab | 1 |

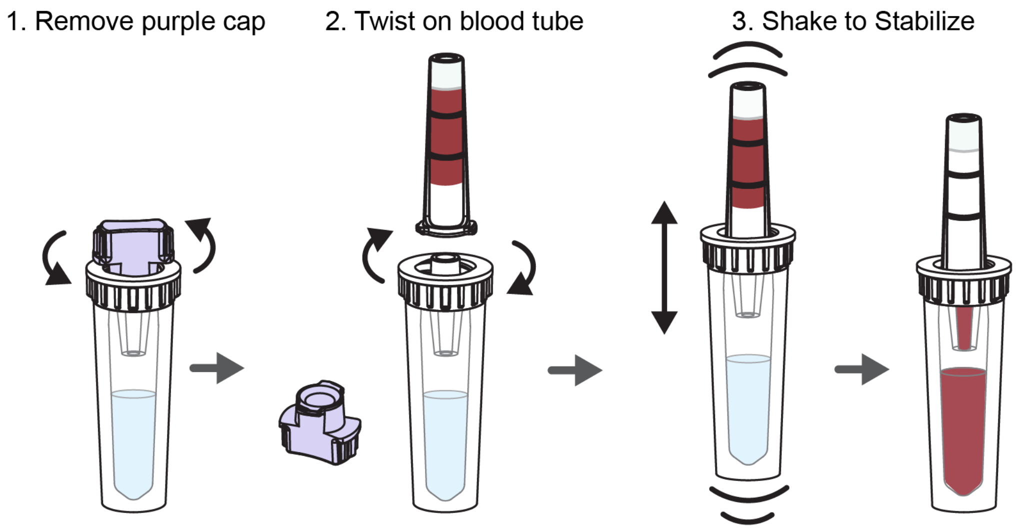


**Figure S2. Schematic workflow of the stabilization process.** The steps for stabilizing a blood sample collected with the Tasso-SST™ are as follows: 1) removal of the purple cap, 2) twisting on the Tasso-SST™ blood tube with the blood sample, 3) shaking up and down vigorously to mix the sample with the stabilizer resulting in RNA stabilized blood.

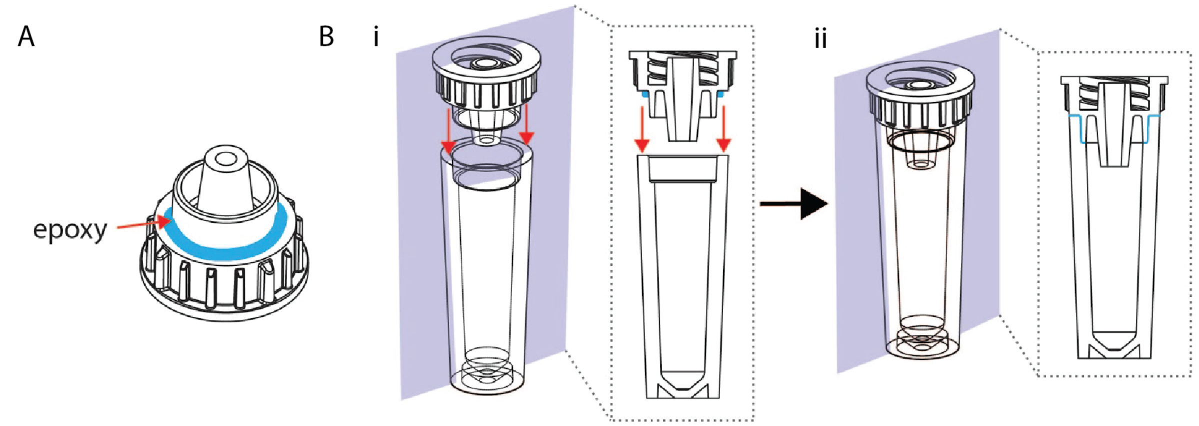


**Figure S3. Fabrication of stabilizer tube.** A) Location where UV epoxy (blue) is applied to the adaptor piece. B) Mating of the adaptor piece with the vial piece showing location of glue before (i) and after (ii) inserting the adaptor piece into the vial piece. Glue is then cured with a 395-405 UV lamp as described in the materials and methods.

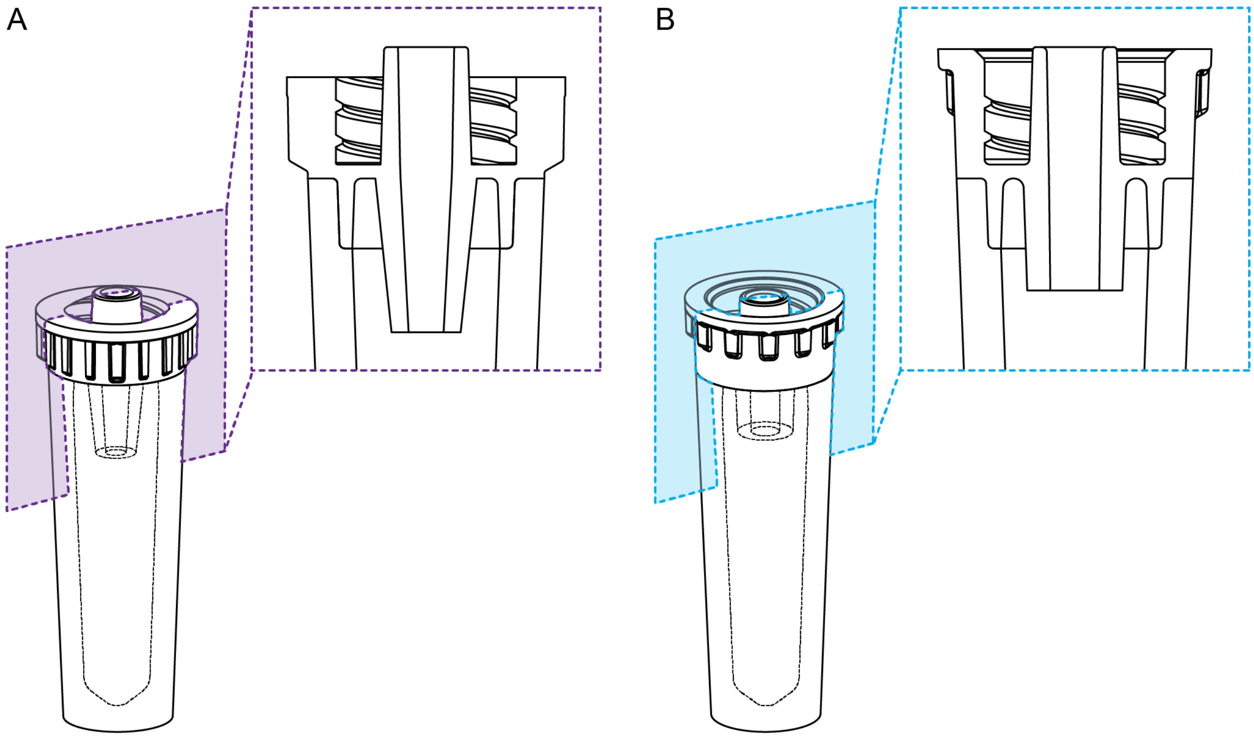


**Figure S4. Tube adaptor design.** A) Tube adaptor design used for groups 1-5. B) Tube adaptor used in groups 6-7. The tube design use for groups six and seven features a wider opening for easier mixing and pipetting.

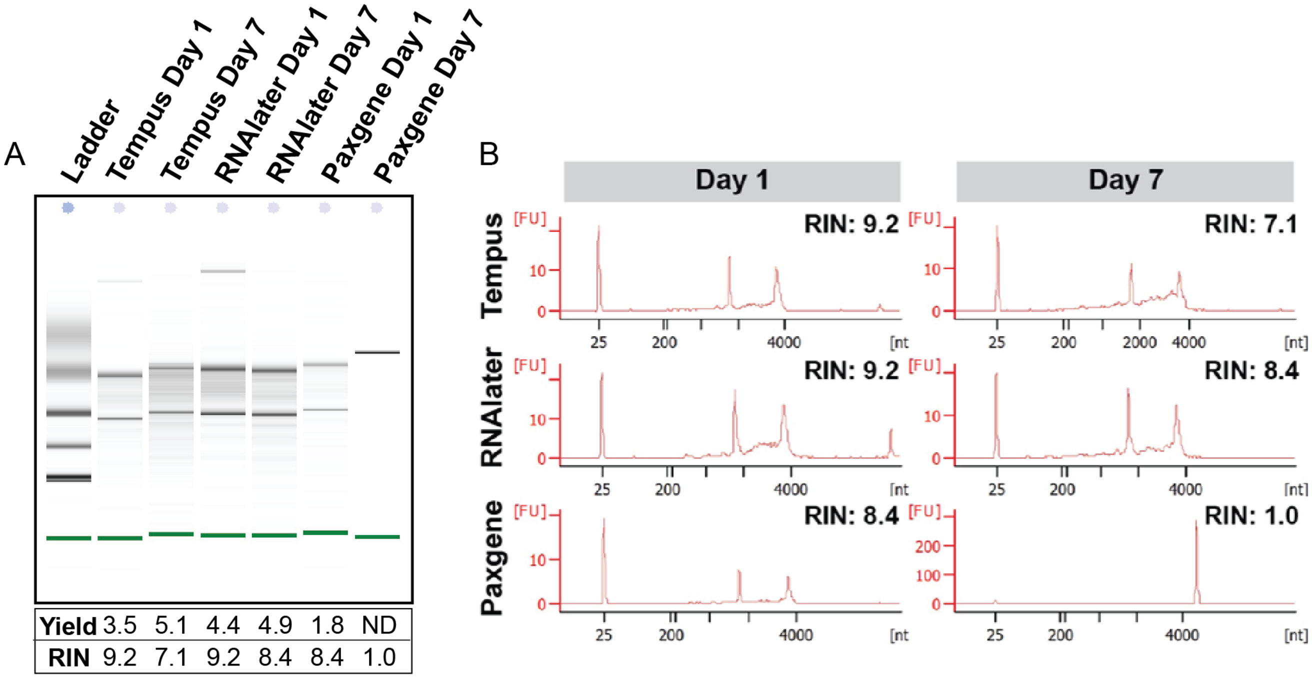


**Fig S5. Performance of select RNA stabilizers on total RNA yield and RNA quality over 7 days at room temperature**. A) Digital gel image, total RNA yield (µg), RIN values and B) electrophoretogram of RNA isolated from blood stabilized in Tempus$^{TM}$, Paxgene® and RNAlater™ for seven days at ambient temperature.

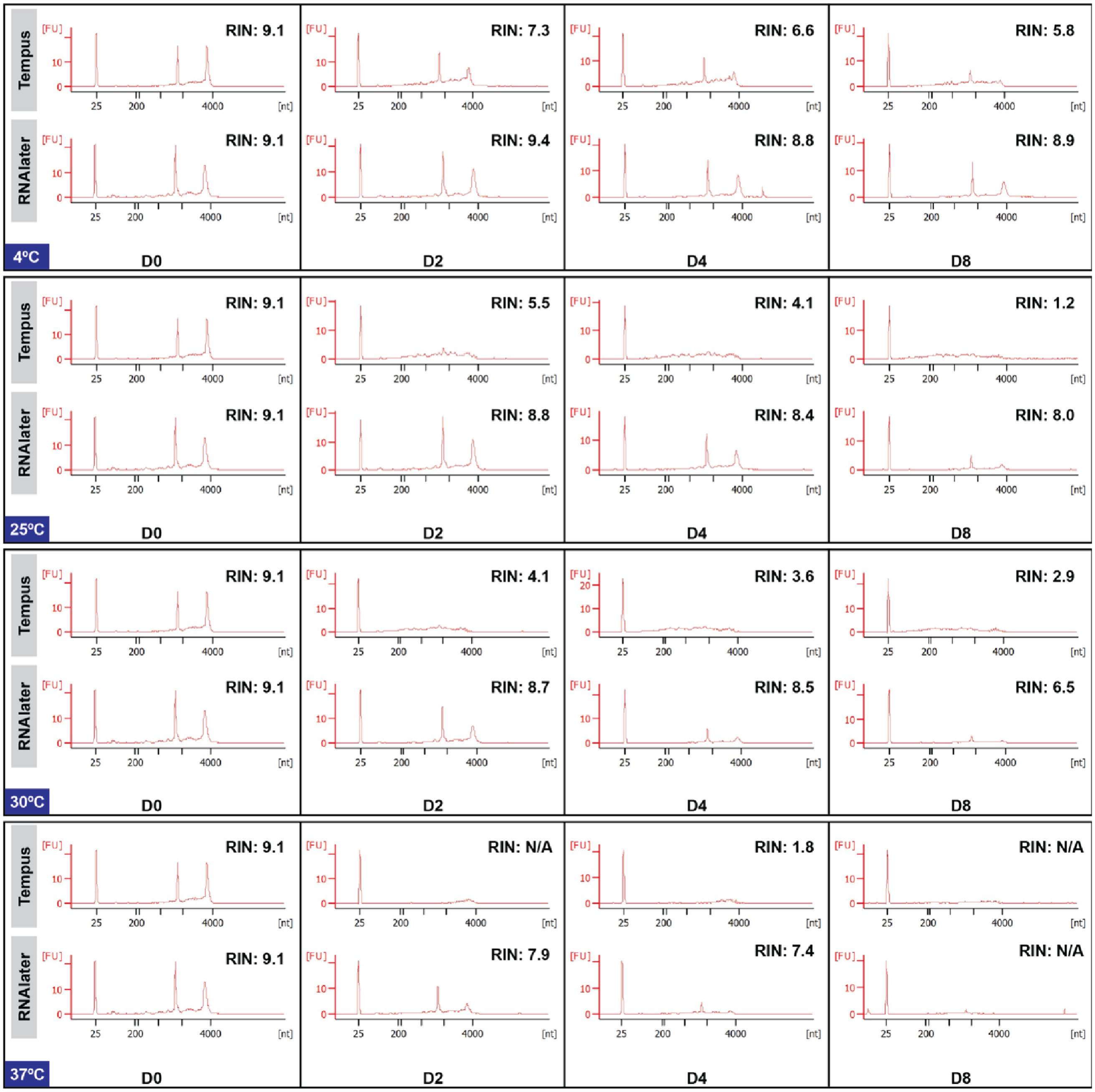


**Figure S6. Electropherogram of RNA samples extracted from blood stored at various temperatures.** Electropherogram of Tempus™ and RNA*later*™ stabilized blood stored at 4°C, 25°C, 30°C and 37°C over 8 days.

**Table S2. Components included in each group**

| Group | IFU version | Video provided | Card with mixing instruction | Tasso-SST™ | Stabilizer Tube | Adaptor 1 or 2 | Hot Pack | Alcohol wipe, bandages | Sample Return bag | Sample Holder |
|---|---|---|---|---|---|---|---|---|---|---|
| 1 | 3.1 | No | No | Yes | Yes | 1 | Yes | Yes | Yes | No |
| 2 | 3.3 | No | No | Yes | Yes | 1 | Yes | Yes | Yes | No |
| 3 | 3.4 | No | No | Yes | Yes | 1 | Yes | Yes | Yes | No |
| 4 | 4.1 | Yes | Yes | Yes | Yes | 1 | Yes | Yes | Yes | Yes |
| 5 | 5.2 | Yes | Yes | Yes | Yes | 1 | Yes | Yes | Yes | Yes |
| 6 | 5.2 | Yes | Yes | Yes | Yes | 2 | Yes | Yes | Yes | Yes |
| 7 | 5.4 | Yes | Yes | Yes | Yes | 2 | Yes | Yes | Yes | Yes |

**Table S3. Logistics for different groups**

| Group | Time of Year Group was sent out | Shipped or Hand Delivered and Picked up | Number of Subjects | Subjects sent one or two kits |
|---|---|---|---|---|
| 1 | Late May, 2020 | Hand delivered and picked up- Seattle Area | 4 | 2 kits |
| 2 | Mid June, 2020 | Shipped- Seattle Area | 5 | 2 kits |
| 3 | Mid July, 2020 | Shipped -Seattle Area | 4 | 1 kit |
| 4 | Mid August, 2020 | Shipped- Nationally (WA, NY, PA, IN) | 13 | 1 kit |
| 5 | Mid September, 2020 | Shipped- Nationally (WA, CO, CA, NE, ME, WI, MA) | 16 | 1 kit |
| 6 | Early November 2020 | Shipped – Seattle Area | 5 | 1 kit |
| 7 | Early December 2020 | Shipped – Nationally (WA, CA) | 5 | 2 kits |

**Table S4. Primer sequences used in gene expression analysis.**

| Gene | Description | NCBI Ref Seq | Primer Sequence (For and Rev) | $T_m$ |
|---|---|---|---|---|
| *GAPDH* | Glyceraldehyde – 3 – phosphate dehydrogenase | NM_002046.7 | 5'-TGCACCACCAACTGCTTAGC-3' | 58ºC |
| | | | 5'- GGCATGGACTGTGGTCATGAG-3' | 58ºC |
| *UBC* | Ubiquitin C | NM_021009.7 | 5'-ATTTGGGTCGCAGTTCTTG-3' | 56ºC |
| | | | 5'-TGCCTTGACATTCTCGATGGT-3' | 56ºC |

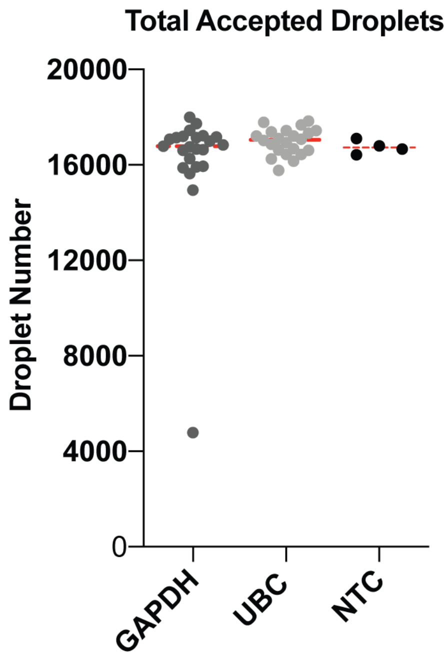


**Figure S7. Total Accepted Droplets of *GAPDH* and *UBC* ddPCR reactions.** NTC represents no template control.

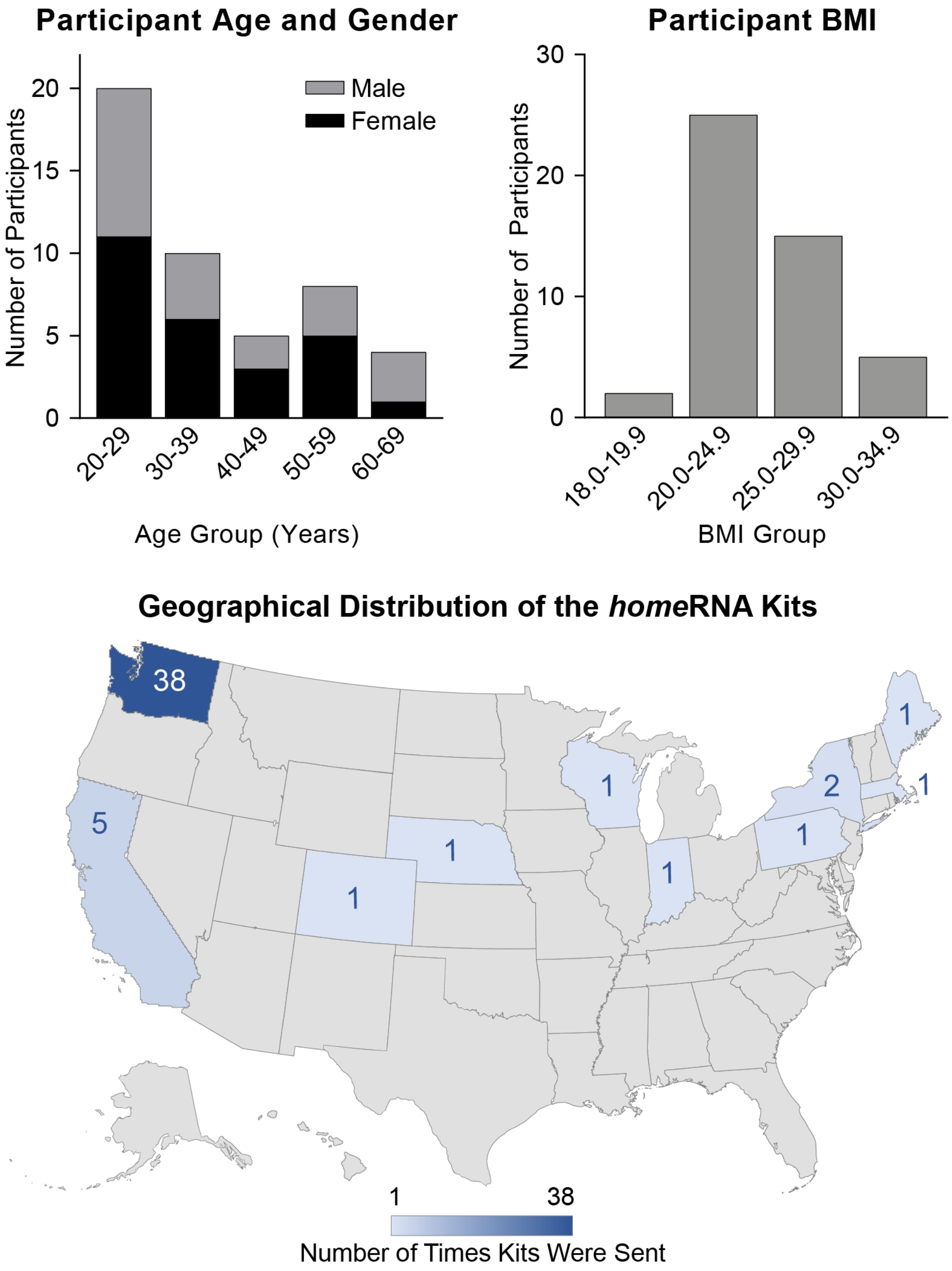


**Figure S8. Age, gender, BMI and region of participants.** A) Age and gender distribution of participants. B) BMI distribuiton of participants. C) Location by state where *home*RNA kits were sent.

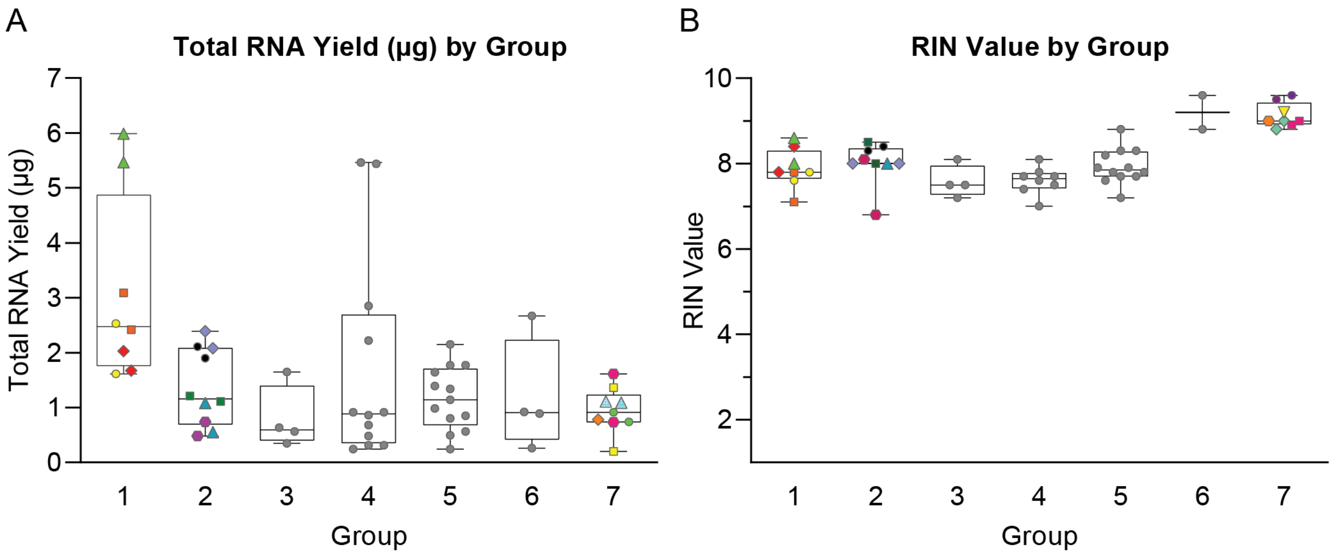


**Figure S9. RNA quantity and quality parameters by participant group.** A) Total RNA yield and B) RIN value by participant group. Grey circles represent a single sample from a single participant. For group 1 and 2 and 7, each participant collected two samples, which are both plotted and represented by a different color and shape for each participant. Box and whisker plots represent full range, interquartile range, and median.

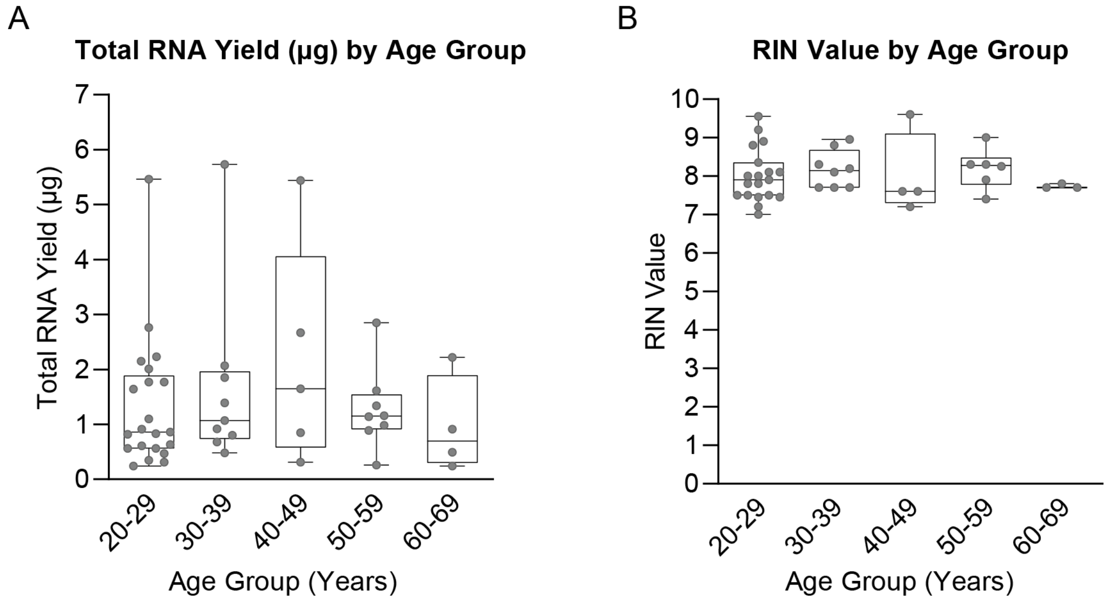


**Figure S10. RNA quantity and quality parameters by age group**. A) Total RNA yield and B) RIN value by age group. The samples taken where each participant collected and stabilized two samples were averaged, and the mean is plotted here. Box and whisker plots represent full range, interquartile range, and median.

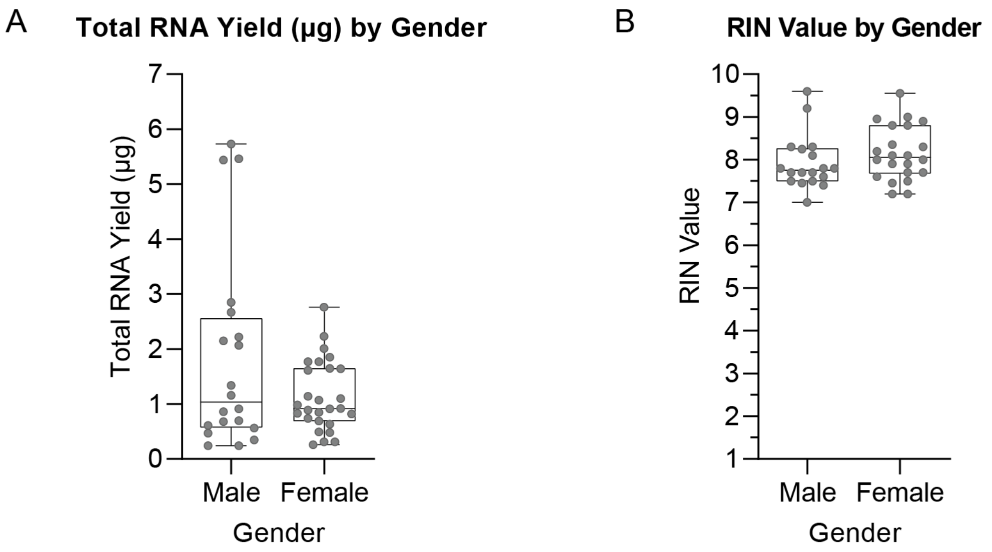


**Figure S11. RNA quantity and quality parameters by gender.** A) Total RNA yield and B) RIN value by gender. The samples taken where each participant collected and stabilized two samples were averaged, and the mean is plotted here. Box and whisker plots represent full range, interquartile range, and median.

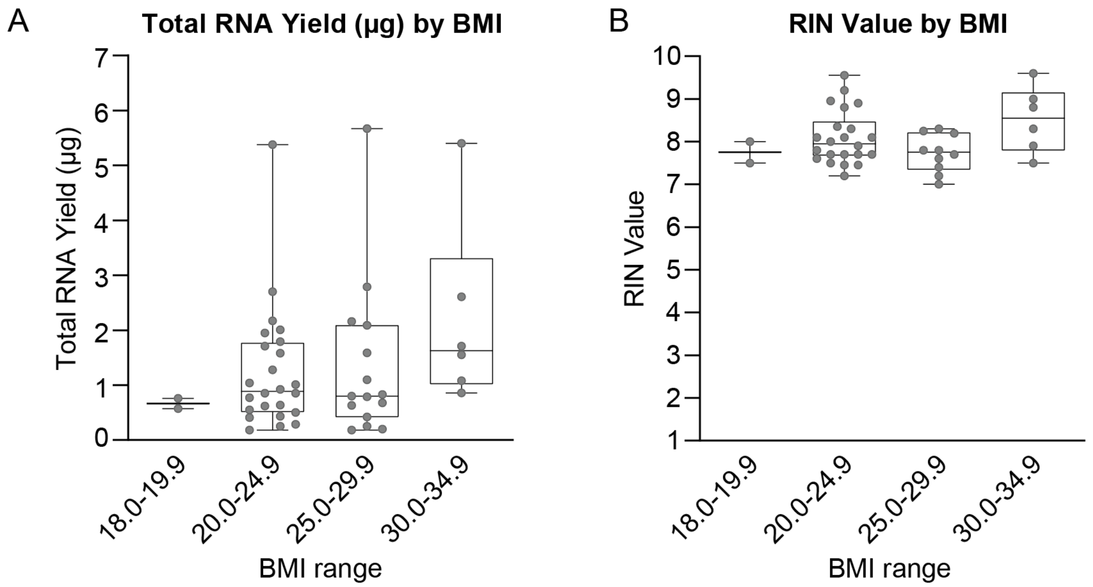


**Figure S12. RNA quantity and quality parameters by BMI.** A) Total RNA yield and B) RIN value by BMI. The samples taken where each participant collected and stabilized two samples were averaged, and the mean is plotted here. Box and whisker plots represent full range, interquartile range, and median.

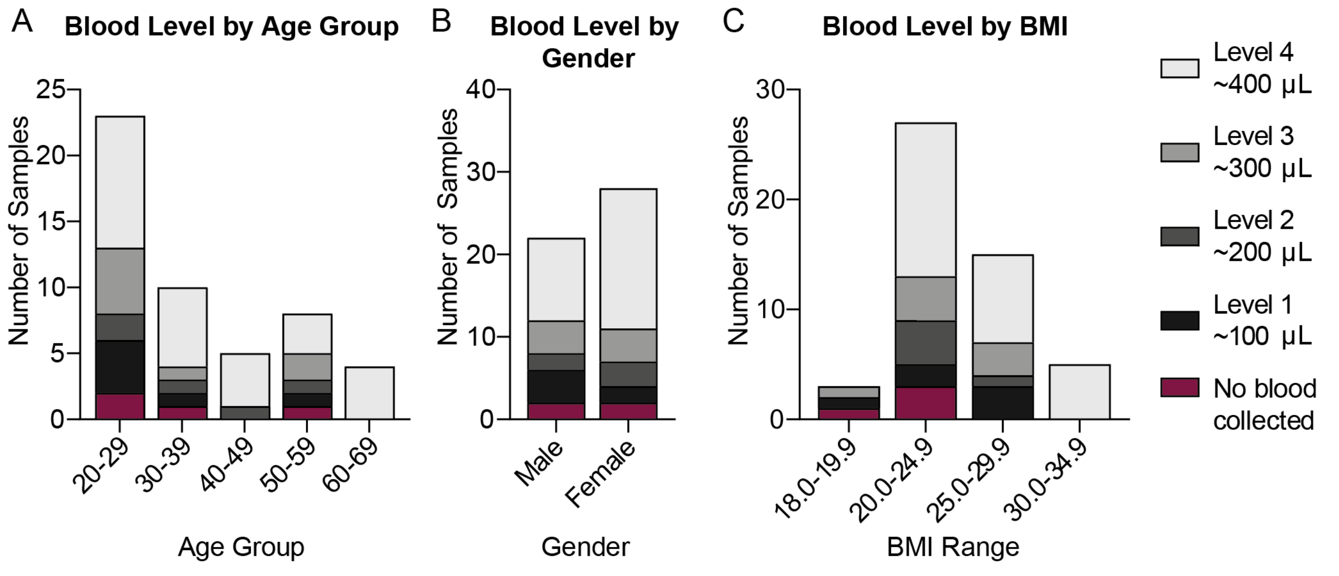


**Figure S13. Reported blood level by participant demographics.** Reported blood level by A) age B) gender and C) BMI of participants.

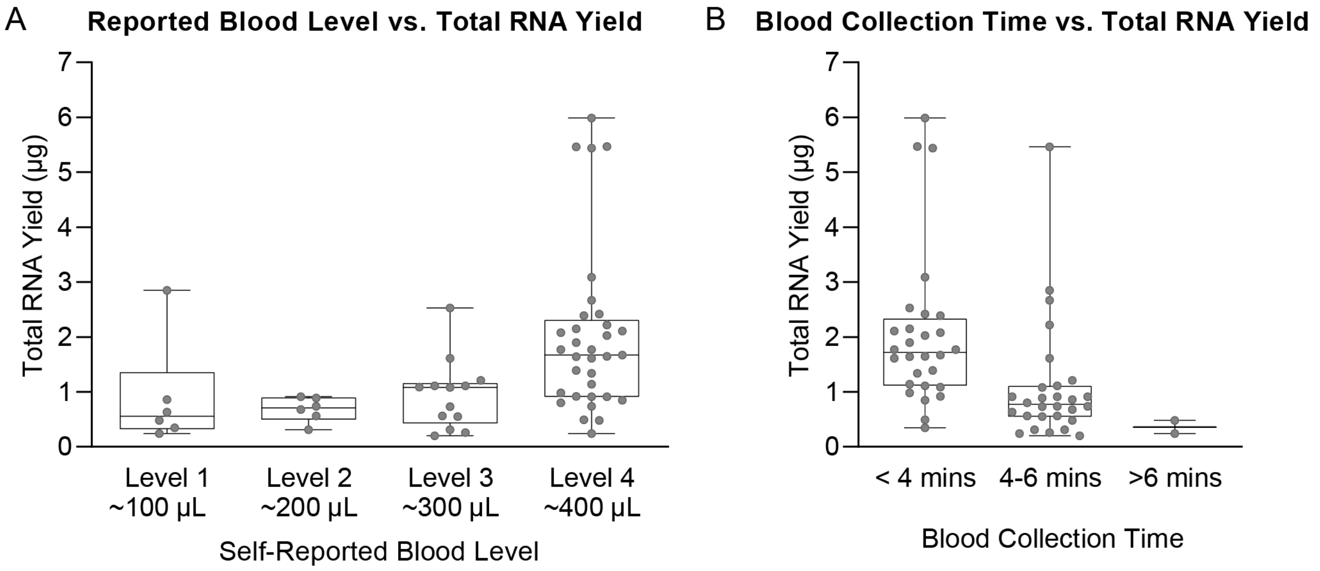


**Figure S14. Total RNA yield in relation to responses to surveys on collection parameters.** A) Reported blood level vs. total RNA yield. B) Blood collection time vs. total RNA yield, where time includes the total time the Tasso-SST™ device was left on the arm. Box and whisker plots represent full range, interquartile range, and median.

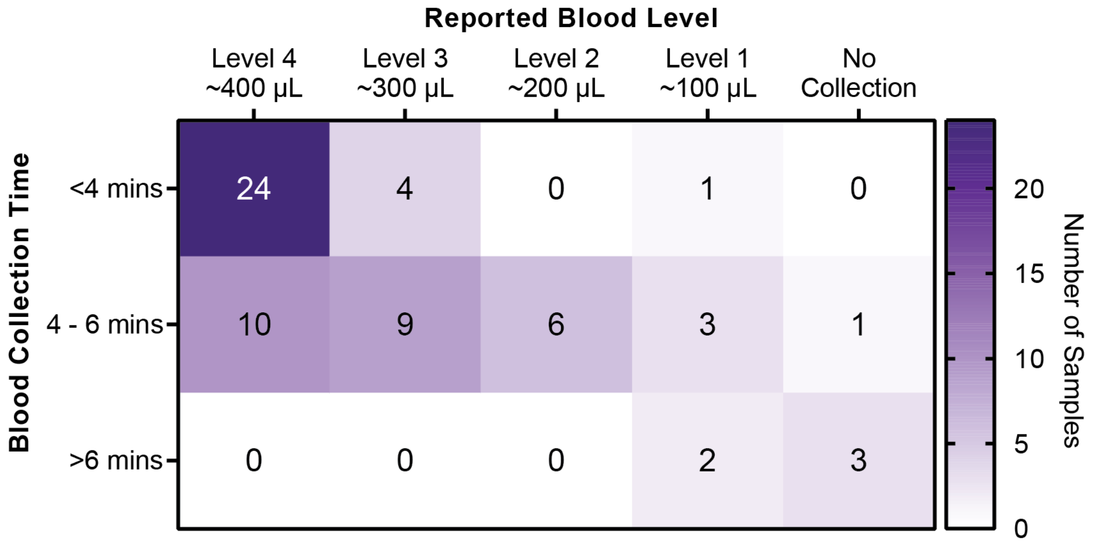


**Figure S15. Reported blood level vs blood collection time.**

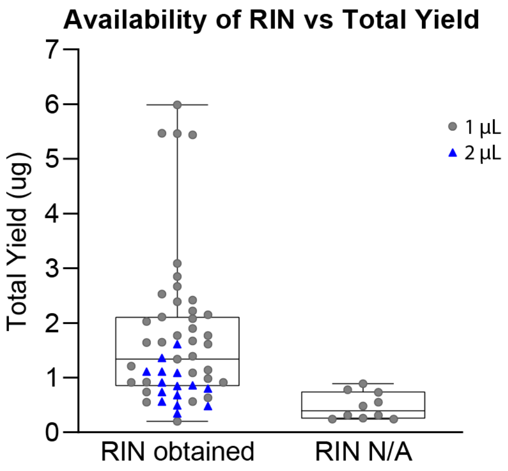


**Figure S16. Availability of RIN based on total yield.** Blue triangles represent samples where 2 µL of isolated RNA were added to the bioanalyzer chip. Grey circles represent where 1 µL was added to the bioanalyzer chip.

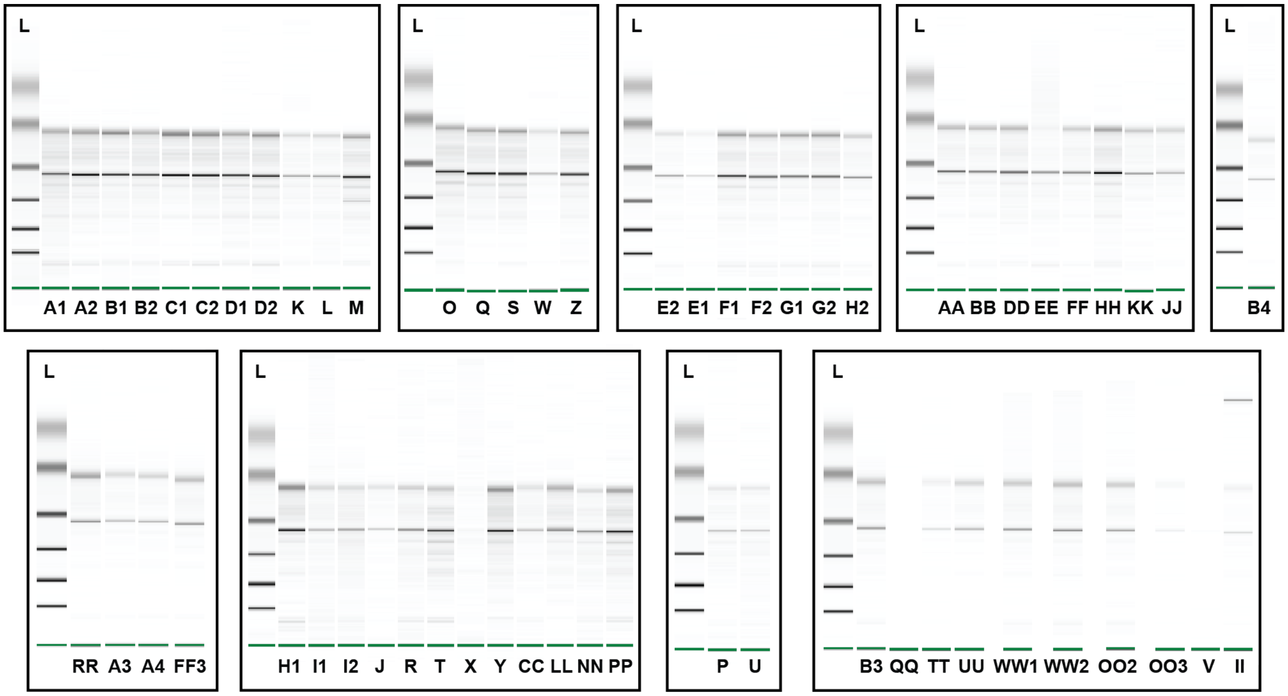


**Figure S17. Digital gel images of all samples obtained from the Bioanalzyer 2100.** Additional sample information corresponding to the letter code are located in the supplemental dataset.

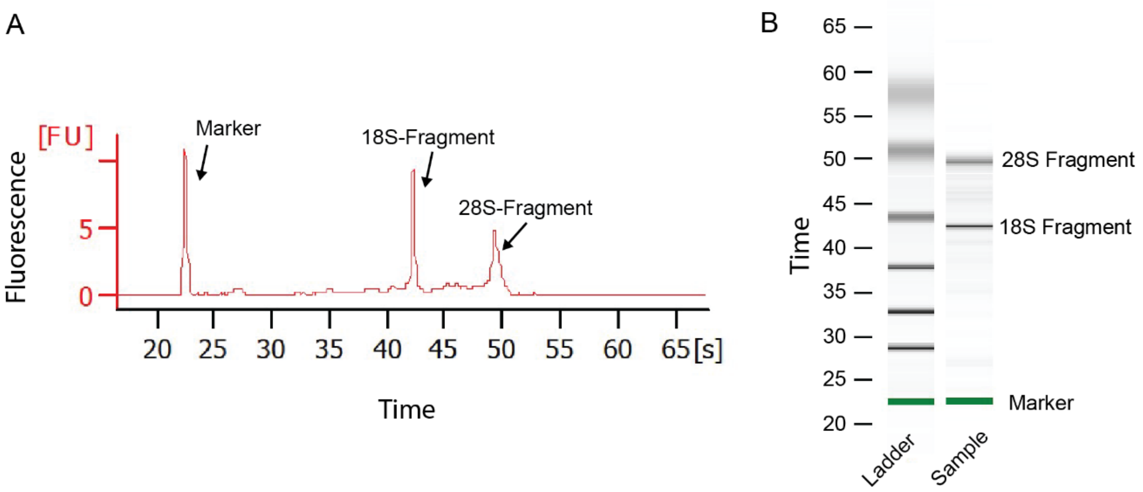


**Figure S18. General Information on Bioanalyzer Profile.** A) Electrophoretogram and B) Digital gel electrophoresis image obtained from Sample B1 annotated with the marker and two other major peaks that help determine RIN including the 18-S fragment peak, and the 28-S fragment peak. More information on interpreting bioanalyzer data, including examples of electrophoretograms obtained from samples with various RIN values can be found in Schroeder 2006.[1]

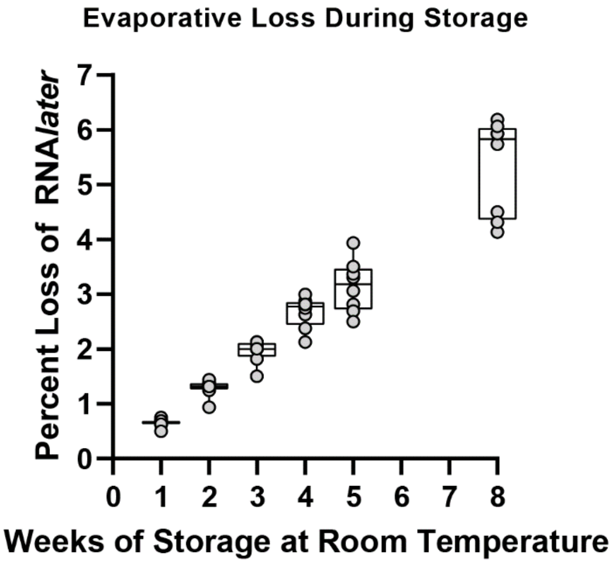


**Figure S19. Evaporative Loss During Storage.** Percentage loss by weight of RNA*later*™ at 1, 2, 3, 4, 5, and 8 weeks storage in the *home*RNA tube at room temperature. Box and whisker plots represent full range, interquartile range, and median.

#### PREPARE DEVICES AND APPLICATION SITE

**FIRST TIME USERS**

Point your phone's camera here to watch an instructional video:

or go to depts.washington.edu/bcmelab/pilotstudy/

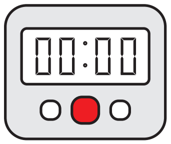


**1. Wash hands and get a timer.**
You will use a timer in Steps 2 and 10.

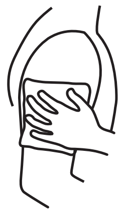

**2. Apply hot pack to upper arm for 2 minutes.**
Warming helps your blood flow better.

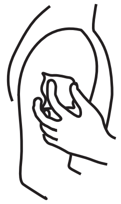

**3. Clean arm with alcohol wipe.**

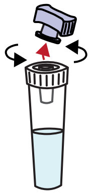

**4. Open the stabilizer tube by twisting off the purple cap.**

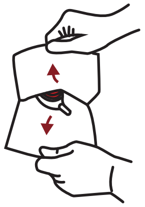

**5. Open Tasso pouch by pulling apart white and clear layers.**
Discard cap in pouch.

#### COLLECT BLOOD USING TASSO-SST

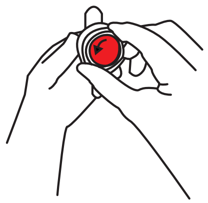

**6. Remove clear plastic cover over the red button.**

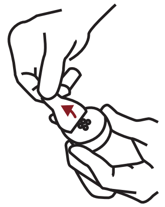

**7. Peel paper tab behind the red button.**
Keep the tube pointing down.

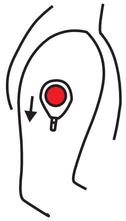

**8. Stick device to shoulder.**
Do not remove once it is on.

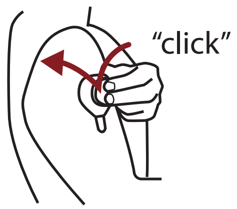


**9. Press button quickly and firmly until it can't go any farther. Wait 2 seconds then let go.**

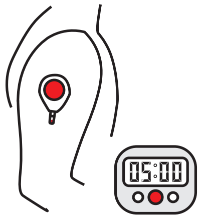


**10. Start a 5 minute timer.**
Keep arm at your side. You wont see blood right away. It can take up to a minute for blood to flow.

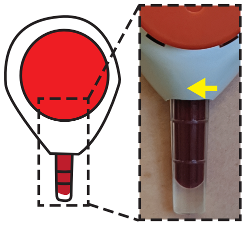

**11. After 5 minutes or when the tube fills, *whichever comes first*, peel off the device.**
It is important to do steps 12-14 as quickly as possible.

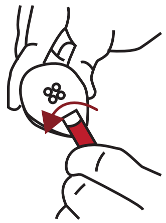

**12. Remove tube by firmly twisting a quarter turn and pulling down.**
This may take a bit of finger strength.

#### MIX AND PACKAGE

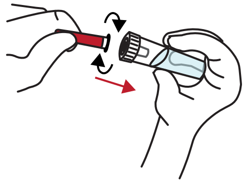

**13. Bring together the blood tube and stabilizer tube and screw these together tightly.**

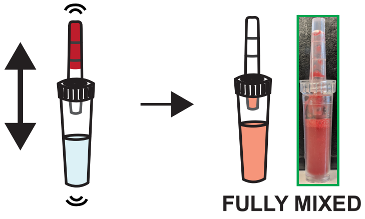


**14. With the stabilizer tube on the bottom, shake hard up and down to mix. Stop when mixed.**
Some fluid may remain in blood tube. When mixed, the color is the same. See insert for details.

DO NOT DISCONNECT TUBES.

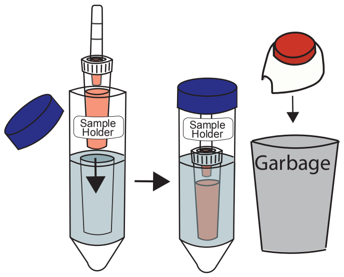


**15. Place sample in sample holder. Throw away used Tasso device.**

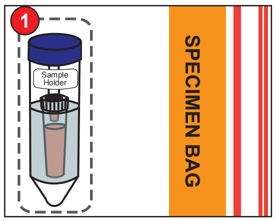


**16. Place blood sample in the specimen bag.**
Leave the absorbent pad in the bag.

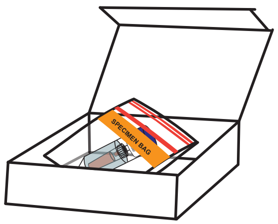


**17. Place specimen bag in box and fill out collection date and time on the box.**
Do not put box in mailer bag until you fill out the online suvey (step **18**).

BCME Pilot 1
[unique code]

**18. Fill out the online survey following the link that was emailed to you.**
You will need unique code and temperature reading located on the box. ***After*** filling out survey, place box in mailer bag and return.

Version 5.6

For research use only

### INSTRUCTIONS FOR USE

#### Home Blood Collection Kit

If you need assistance or have questions regarding the use of this kit, contact us at [redacted] or call us at [redacted]

**Thank you for participating in our study!**

#### KIT CONTENTS

Make sure your kit contains all components listed below

**1) Tasso-SST device**

**2) Stabilizer tube**

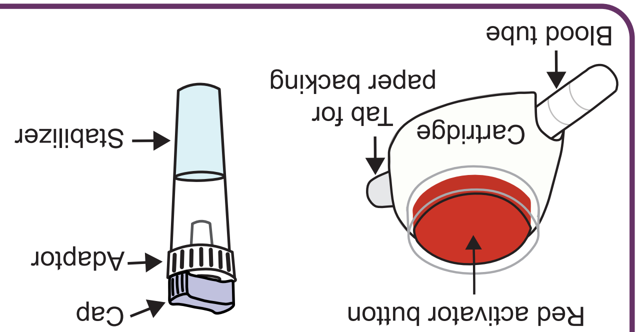


**3) Alcohol wipes**

**4) Hot pack**

**5) Bandage**

**6) Specimen bag**

**7) Sample Holder**

**8) Mail return bag**

#### INTENDED USE

TASSO-SST is a single use blood collection device that is intended for the self collection of capillary blood from the upper arm of adults (18 years or older). The stabilizer tube contains liquid that is intended for stabilizing the collected blood. The collected sample is mailed back to the BCME research laboratory for academic research use.

#### STORAGE

Store at 15 - 30ºC (60 - 80ºF) in a dry place.

#### WARNINGS

- The TASSO-SST is a sterile device. Do not open until use.
- The TASSO-SST device contains sharps. Handle with care.
- For external use only.
- Keep out of reach of children.
- Use while seated as fainting may occur during blood sampling procedure.
- Do not activate the red button until device is firmly on skin.
- Wipe application site with alcohol wipe to reduce infection risk.
- Always use a new unopened pouch of Tasso-SST. Do not re-use.
- Discard used Tasso-SST device in regular garbage.

#### ONLINE SURVEY

Please check your email for a link to fill out an online survey regarding your experience on using the home blood collection kit. If you cannot find the link, please email us at [redacted] and we can send you another link.

#### RETURNING KIT

Place box in provided mail return bag and seal. Store sealed bag containing the blood collection kit box at room temperature. Your kit will be picked up from the location you specified within 24 hours. Please notify us if your kit did not get picked up by emailing [redacted] or calling [redacted]
